# Remote contextual fear retrieval engages activity from salience network regions in rats

**DOI:** 10.1101/2021.12.22.473939

**Authors:** Moisés dos Santos Corrêa, Gabriel David Vieira Grisanti, Isabelle Anjos Fernandes Franciscatto, Tatiana Suemi Anglas Tarumoto, Paula Ayako Tiba, Tatiana Lima Ferreira, Raquel Vecchio Fornari

## Abstract

The ability to retrieve contextual fear memories depends on the coordinated activation of a brain-wide circuitry. Transition from recent to remote memories seems to involve the reorganization of this circuitry, a process called systems consolidation that has been associated with time-dependent fear generalization. However, it is unknown whether emotional memories acquired under different stress levels can undergo different systems consolidation processes. Here, we explored the activation pattern and functional connectivity of key brain regions associated with contextual fear conditioning (CFC) retrieval after recent (2 days) or remote (28 days) memory tests performed in rats submitted to strong (1.0mA footshock) or mild (0.3mA footshock) training. We used brain tissue from Wistar rats from a previous study, where we observed that increasing training intensity promotes fear memory generalization over time, possibly due to an increase in corticosterone (CORT) levels during memory consolidation. Analysis of Fos expression across 8 regions of interest (ROIs) allowed us to identify coactivation between them at both timepoints following memory recall. Our results showed that strong CFC elicits higher Fos activation in the anterior insular and prelimbic cortices during remote retrieval, which was positively correlated with freezing along with the basolateral amygdala. Rats trained either with mild or strong CFC showed broad functional connectivity at the recent timepoint whereas only animals submitted to the strong CFC showed a widespread loss of coactivation during remote retrieval. Post-training plasma CORT levels are positively correlated with FOS expression during recent retrieval in strong CFC, but negatively correlated with FOS expression during remote retrieval in mild CFC. Our findings suggest that increasing training intensity results in differential processes of systems consolidation, possibly associated with increased post-training CORT release, and that strong CFC engages activity from the aIC, BLA and PrL – areas associated with the Salience Network in rats – during remote retrieval.

## INTRODUCTION

Traumatic events may induce pathological forms of fear memory, including the late-onset pathological fear generalization observed in patients with post-traumatic stress disorder (PTSD). This symptom is marked by unwanted recurring thoughts of a traumatic event in contexts that are different from those in which the traumatic event originated (Mahan and Ressler, 2012). Understanding which brain regions are engaged during the retrieval of such memories is imperative for the development of new pharmacological and neuromodulatory treatments. Recent advances suggest that long-term contextual fear memory retrieval seems to be dependent on the coordinated activation of a brain-wide engram circuit (Rao-Ruiz et al., 2021; Wheeler et al., 2013). Furthermore, the transition of a recent to remote memory is thought to involve a reorganization of this circuit, a process called systems consolidation (DeNardo et al., 2019; Moscovitch et al., 2016). Traditionally, recent contextual fear conditioning (CFC) retrieval is associated with high contextual specificity and neuronal activity in the hippocampal formation, especially its dorsal portion (dHPC). With time, contextual fear memories become more generalized, less affected by hippocampal lesions, and more associated with activity in neocortical regions (Frankland et al., 2004; Winocur et al., 2010). Nonetheless, evidence also suggests that the dHPC may be engaged in remote retrieval (Barry and Maguire, 2019; Vetere et al., 2021), whereas the medial prefrontal cortex (mPFC) may also be involved in the retrieval of precise remote memories (Matos et al., 2019; Moscarello and Maren, 2018). Hence, the mechanisms by which the temporal reorganization of the engram circuit occurs and how it affects memory generalization are still controversial.

Parallel to the research into systems consolidation and time-dependent fear generalization, other studies also suggest that recognition, spatial or emotional memory engram cells are allocated during acquisition in several brain regions other than the hippocampus (Roy et al., 2022, 2016). Some of these regions are the amygdala (Redondo et al., 2014), the anterior insular cortex (aIC) (Sano et al., 2014) and the mPFC, especially the prelimbic cortex (PL) (Zhang et al., 2011). These regions are part of the so-called Salience Network (SN) (Grandjean et al., 2020) and are highly responsive to arousing situations in humans (Hermans et al., 2014a) and in mice (Mandino et al., 2021; Zerbi et al., 2019). Strong, aversive events are thought to prompt a shift of memory encoding dependence from a hippocampal-dominated engram circuit to SN regions during memory acquisition and consolidation, favoring the consolidation of emotional memories (Arnsten, 2009; Hermans et al., 2014b). In contrast, there is evidence that CFC in rats is associated with decreased functional connectivity between the retrosplenial cortex (RSC) – a region considered part of the “Default-mode-like network” (DMN-like, to distinguish it from the human DMN) in rodents – to the aIC (Gozzi and Schwarz, 2016; Ji et al., 2018; Upadhyay et al., 2011). Furthermore, a comprehensive neuroimaging study using human data showed that generalized fear is associated with decreased activation in nodes of the DMN, especially the HPC, and increased activation of the aIC, mPFC and ACC (including its dorsal and subgenual portions) (Webler et al., 2021). Hence, this shift in memory systems resulting from higher stress levels and the engagement of different neurocognitive networks elicited by strong aversive situations may trigger different processes of systems consolidation.

Several studies have also linked increasing levels of arousal and stress hormones with strengthened contextual fear memories and to an acceleration of the rate of time-dependent fear generalization and systems consolidation (Dos Santos Corrêa, et al., 2021; Pedraza et al., 2016; Schwabe et al., 2022). More recently, studies showed that post-training activation of the noradrenergic and glucocorticoid systems modulates contextual fear memory specificity (Atucha et al., 2017; Dos Santos Corrêa et al., 2021; Gazarini et al., 2021; Roozendaal and Mirone, 2020). Therefore, it is possible that interfering with arousal levels by changing training parameters may affect the processes of time-dependent fear generalization and systems consolidation through changes in the engagement of neurocognitive networks (Schwabe, 2017; Zhang et al., 2022). To explore this question, a recent study from our laboratory investigated the rate of time-dependent fear memory generalization following increasing intensities of CFC training. Our data showed that mild footshocks (0.3mA) elicited lower corticosterone (CORT) levels and accurate remote contextual memories while, on the other hand, strong footshocks (1.0mA) increased CORT release during memory consolidation, which was associated with remote generalized fear (Dos Santos Corrêa et al., 2019). These results indicate that increasing levels of arousal elicited different appraisal of the aversive experience, promoting fear memory generalization possibly due to an increase in glucocorticoid activity during memory consolidation.

In the present study we examined the expression of the immediate early gene product Fos following memory retrieval in the same rats used in the previous study (Dos Santos Corrêa et al., 2019). Here, we explored whether there were changes in functional connectivity of key brain regions associated with the SN, the DMN-like or both networks (Mandino et al., 2021; Zerbi et al., 2019). We hypothesized that increasing footshock intensity during CFC training would elicit different processes of systems consolidation, which can be observed through changes in functional connectivity between these regions during recent and remote retrieval tests.

## METHODS

### 1 Subjects, behavioral task, and post-training plasma CORT quantification

For the present study, we used a subset of animals from a previously published experiment (Dos Santos Corrêa et al., 2019) and the behavioral data from these animals was extracted from the previously published dataset ([dataset] Dos Santos Corrêa et al., 2019). These animals were three-month-old male Wistar rats, obtained from *Instituto Nacional de Farmacologia* (INFAR-UNIFESP (total n = 42; weighing between 275-385g at the time of training). All rats were kept in controlled conditions of temperature (23 ± 2°C) and maintained on a 12 h light/dark cycle (light phase starting at 7 am) with free access to food and water. Behavioral experiments were conducted during the light phase of the cycle, from 11 am to 4 pm, during the rat’s nadir of corticosterone circadian rhythm. All procedures were conducted according to the guidelines and standards of CONCEA - *Conselho Nacional de Controle de Experimentação Animal* (Brazilian Council of Animal Experimentation) and were previously approved by the Ethics Committee on Animal Use - UFABC (CEUA - protocol numbers 5676291015 and 7479070916)

Contextual fear conditioning experiments were conducted as described before (Dos Santos Corrêa et al., 2019) in a windowless room containing the conditioning chamber (32 cm wide, 25 cm high and 25 cm deep, Med-Associates VFC-008). Rat freezing behavior was monitored via a frontal near infra-red-light camera. Freezing was assessed using an automated scoring system (Video Freeze, Version 1.12.0.0, Med-Associates), which digitized the video signal at 30 Hz and compared frame by frame movement to determine the amount of time spent freezing. The context was characterized by a grid floor composed of 20 stainless steel rods (diameter: 4.8 mm), top and front walls made of transparent polycarbonate, a back wall made of white acrylic, stainless-steel sidewalls and a drop pan below the floor grid. The light in the conditioning box remained on, and background noise was emitted during the training and test sessions. The chamber was cleaned with alcohol 10% before and after each rat.

In all experiments, rats from each home-cage were randomly assigned to one of the experimental or control groups, and all groups were matched according to the average body weight. Animals were trained and tested in batches, with a balance of experimental groups but in separate experiments for each timepoint. During training, rats were placed in the conditioning chamber for four minutes. After two minutes they were presented with three unsignalled footshocks (1 s duration, 0.3 or 1.0 mA, 30 seconds apart). Following the last footshock rats remained in the chamber for another minute, and then were returned to their home cage. Control rats underwent the same training procedure but did not receive the footshocks during the conditioning session (Context Only – C.O.). Thirty minutes after each training session, blood was collected from the tail of each rat and post-training plasma CORT levels were quantified by ELISA according to the described in Dos Santos Corrêa et al. (2019). CORT data are also published in ([dataset] Dos Santos Corrêa et al., 2019). After each training condition, separate groups of rats were tested either 2 days (C.O.|Recent, n = 5; 0.3mA|Recent, n = 6; 1.0mA|Recent, n = 6) or 28 days (C.O.|Remote, n = 5; 0.3mA|Remote, n = 6, 1.0mA|Remote, n = 6) later. During testing, rats were placed back in the training context for five minutes and freezing time was assessed. No footshock was delivered during test sessions. Another control group of rats was only handled but did not undergo CFC training or testing (Homecage, Recent, n = 4, Remote, n = 4).

### 2 Perfusion and histology

Ninety minutes following the end of the test session, rats were deeply anesthetized with 30% urethane and then perfused transcardially with 100 mL saline followed by 500 mL of ice-cold 4% paraformaldehyde (PFA) dissolved in phosphate buffer saline. The brains were removed, fixed for at least 2 hours in PFA, then transferred to 30% sucrose solution and stored at 4°C. After dehydration, these brains were frozen in isopentane on dry ice and stored at -80 °C. Frozen coronal sections (40 µm) were cut in a microtome with dry ice and stored in 5 serial sets. The second set was collected in glass slides and Nissl-stained for cytoarchitectural delineation of brain areas, and the first and eventually the third sets were used for Fos immunolabeling.

### 3 Immunohistochemistry and cell counting

Free-floating sections underwent antigen retrieval (10mM Sodium Citrate, ph 8.5) for 30 minutes at 80 ºC, and peroxidase blocking (0.3% hydrogen peroxide in 0.02M PBS) followed by indirect antibody staining protocols aiming Fos labeling by avidin-biotin reaction. Each immunohistochemistry assay included sections from all experimental groups from both timepoints (including homecage animals) to avoid a statistical artefact due to inter-assay variability. Sections were incubated at 4 ºC for 72 hours with rabbit anti-cfos polyclonal antibody (1:10,000 AbCam) in 2% normal goat serum, 0.3% Triton X-100 and 0.02M PBS. After washes, incubation with secondary antibody (anti-rabbit Ab from goat diluted in 1:200 – BA1000, Vector, in 0.02M PBS and 0.3% Triton X-100) took 90 minutes at room temperature. The secondary antibody was biotinylated in the avidin-biotin-complex solution (1:200, ABC Kit, VectaStain Elite, Vector) in 0.02M PBS and 0.3% Triton X-100. The peroxidase complex was visualized using the chromogen diaminobenzidine 3,3-tetrachloride (DAB Kit, Vector). Finally, sections were mounted on gel-coated slides and left to dry for at least 48 h. After drying, sections were diaphanized, coverslipped with DPX mountant medium (Sigma 06522) and left to dry for at least a week.

Fos expression was analyzed in 8 regions of interest (ROIs, Table 1). Images from each ROI were acquired using either a Zeiss microscope model Axio image 2 at 20x magnification or a Leica microscope model DM5500 at 10x magnification, following the same frame size, image size and area for each microscope. One ROI (aRSC) was smaller than the standard area and was delimited using the ImageJ software (NIH, Washington, United States). The anatomical delimitation of each ROI was based on Paxinos and Watson rat brain atlas (Paxinos and Watson, 2007) and adjacent Nissl-stained sections. Table 1 shows the information regarding anteroposterior coordinates of slices, image size, amplification and number of bilateral sections for each ROI. ImageJ®L (1.52a) was used to count cells automatically following parameters of size, circular shape, and contrast level adapted to each image but within a pre-established range to ensure consistency. Experimenters were blinded to the experimental group. The cell density of each photo was obtained by dividing the number of cells by the total area of the image. Mean Fos density of the photos from each animal, either for recent or remote timepoints, was normalized by the averaged Fos density of homecage rats that were euthanized on the same days as the other groups. The experimental unit was the cell Fos density normalized by the homecage group for each animal. One case from the 0.3mA|Remote group was discarded due to issues after multiple immunohistochemical essays, lowering the number of cases in this group down to 5. Counts from sections with tissue damage were excluded from the analysis.

**Table 1.** Description of microscopy for each region of interest (ROI)

| Region | Microscope | Bregma Range | Number of<br>bilateral frames<br>per ROI | Size (pixels)<br>per photo | µm per pixel | Lens | Median<br>bilateral<br>sections per<br>animal |
| --- | --- | --- | --- | --- | --- | --- | --- |
| **Hippocampus - Dentate Gyrus (DG)** | Zeiss | [-2.28,-3.72] | 2 | 1388:1040 | 0,5 | 20x | 2 |
| **Hippocampus - CA3** | Zeiss | [-2.28,-3.72] | 1 | 1388:1040 | 0,5 | 20x | 2 |
| **Hippocampus - CA1** | Zeiss | [-2.28,-3.72] | 3 | 1388:1040 | 0,5 | 20x | 2 |
| **Prelimbic (PrL)** | Leica | [4.2,2.52] | 1 | 3072:2204 | 0,3 | 10x | 3 |
| **Basolateral Amygdala (BLA)** | Zeiss | [-1.56,-3.24] | 1 | 1388:1040 | 0,5 | 20x | 2 |
| **Anterior Retrosplenial Cortex (aRSC)** | Leica | [-2.28,-3.72] | 1 | 2410:1807 | 0,3 | 10x | 3 |
| **Anterior Cingulate Cortex (ACC)** | Zeiss | [3.24,2.28] | 1 | 1388:1040 | 0,5 | 20x | 2 |
| **Anterior Insular Cortex (aIC)** | Zeiss | [3.24,0.72] | 1 | 1388:1040 | 0,5 | 20x | 3 |

Note: The frames were taken from an interval of bregma coordinates that ranged from the most anterior to the most posterior for each brain region (column “Bregma”). Brain regions larger than the microscope’s standard area were obtained in more than one frame (column “Number of bilateral frames per ROI”). Each animal had at least 1 and up to 3 bilateral brain sections to capture frames from; the column “Median bilateral sections per animal” shows the median number of bilateral sections per animal taken from all experimental groups in both recent and remote timepoints.

### 4 Statistical Analysis

#### 4.1 Behavior and Fos density for each ROI

All descriptive and inferential statistical analyses for behavior and Fos density were done using JAMOVI (version 1.8) and Python. The conditioned fear response to context was quantified as the percent time the animal spent freezing during re-exposure to the training context. Behavioral results are expressed as the group mean percent freezing time ± standard error of the mean (S.E.M). Immediate early gene activation results are expressed as the group mean normalized Fos density ± S.E.M. All data were checked for the assumptions of normality and homogeneity of variances with the tests of Kolmogorov-Smirnov and Levene.

Behavioral data and Fos differences for each brain region were analyzed using one and two-way ANOVAs, respectively, with the between-subject factors being Footshock and Timepoint. The Tukey post hoc test was further used to identify significant differences when applicable. Significance for all tests described in this section was set at p ≤ 0.05. Effect sizes for ANOVAs (*η*^2^p) and post hoc tests (Cohen’d) are reported only when the test was found significant (“*η*^2^p” values above 0.14 are considered large effects; values between 0.06 and 0.14 are considered moderate; and below 0.06, small) or in the Supplementary Material. Associations between freezing times and Fos density for each region were calculated using bicaudal Pearson correlations (totaling 16 correlations, uncorrected for multiple comparisons), including data from all groups exposed to the context in each timepoint. Each set of correlations was calculated from a vector of size 15-17, and was displayed as color-coded correlation matrices using Python.

#### 4.2 Fos Coactivation analysis

In experimental animal studies, neuronal activity is usually measured by evaluating changes in the expression of immediate early genes such as *c-fos* or their products. Functional connectivity for these measures is assessed by calculating covariance across subjects, rather than within-subjects (Ben-Ami Bartal et al., 2021; Careaga et al., 2019; Coelho et al., 2018; Worley et al., 2020). Within each of the six experimental groups of animals (retrieval test at recent or remote timepoints; Context Only, 0.3 and 1.0 mA), all possible pairwise correlations between the Fos signal within the eight brain regions analyzed were determined by computing Spearman correlation coefficients (totaling 192 correlations, uncorrected for multiple comparisons). Each set of correlations was calculated from a vector of size 5-6, and was displayed as color-coded correlation matrices using Python. Fos coactivation (i.e., functional connectivity) was considered when the Spearman’s rank correlation coefficient was below a threshold of the one-tailed significance level of *p* ≤ 0.025 and *rho* ≥ 0.81. To determine how Fos coactivation correlations differed by footshock intensity and timepoint, we contrasted mean r values between all ROIs for each group using a one-way Welch ANOVA. Games-Howell post hoc tests were used for multiple comparisons between groups. Significance for the ANOVA and post hoc tests described in this section was set at *p* ≤ *0*.*05*. To investigate the effects of footshock intensity and timepoint on individual ROI by ROI correlations (i.e., is PrL more strongly correlated with BLA at the Remote timepoint versus Recent for each footshock intensity, or at Mild versus Strong training for each timepoint), we contrasted the correlation of each condition to the others (all possible comparisons) using the Fisher r to z transformation and unicaudal z test to determine *p* values for each contrast. The set of p-values was then globally adjusted to correct for multiple comparisons using Benjamini and Hochberg’s false discovery rate procedure (Worley et al., 2020) maintaining a false discovery rate of 25%. Pairwise ROI by ROI comparisons identified as significant using this method are summarized in Table 3.

#### 4.3 CORT levels and Fos density for each ROI

We performed correlation analyses between post-training CORT levels and Fos expression from the eight brain regions analyzed. Within each of the six experimental groups of animals, correlations were determined by computing bicaudal Spearman correlation coefficients (totaling 48 correlations, uncorrected for multiple comparisons). Each set of correlations was calculated from a vector of size 5-6, and was displayed as color-coded correlation matrices using Python. To determine how Fos and CORT correlations differed by footshock intensity and timepoint, we contrasted averaged r values between all ROIs for each group using a one-way Welch ANOVA. Games-Howell post hoc tests were used for multiple comparisons between groups. The level of significance for the correlation, Welch ANOVA and post hoc tests described in this section was set at *p* ≤ 0.05.

## RESULTS

### 1. Remote memory retrieval is associated with higher neocortical functional connectivity

The brains of a subset of rats used in a previous study (Dos Santos Corrêa et al., 2019) were used for analyses of the *c-fos* activity-dependent neuronal expression. The behavioral data of this subset of animals is representative of the main results previously published (Figure 1a). One-way ANOVA indicated that increasing footshock intensity elicited higher freezing times in both recent [Footshock (F(2,14) = 53.0, *p* < ◻.001, *η*^*2*^*p* = 0.88)] and remote timepoints [Footshock (F(2,13) = 24.60, *p* <◻ .001, *η*^*2*^*p* = 0.79)]. In both timepoints, rats from the C.O. groups showed significantly lower freezing when compared to the 0.3 (Recent, *p* = 0.01; Remote, *p* = 0.02) and 1.0 mA groups (Recent and Remote, *p* <◻.001). In addition, rats from the 0.3 mA group showed significantly lower freezing times when compared to the 1.0 mA group in both timepoints (Recent, *p* <◻0.001; Remote, *p* = 0.003, see table S1 for all post hoc results and effect sizes). These results were associated with remote contextual discrimination after low intensity training (0.3 mA) and remote contextual generalization after high intensity training (1.0 mA; Dos Santos Corrêa et al., 2019). Post-training plasma CORT levels for each of the three footshock intensities in this subset of animals and inferential statistics can be seen in the supplementary material Fig S1 e Table S2.

**Figure 1:**
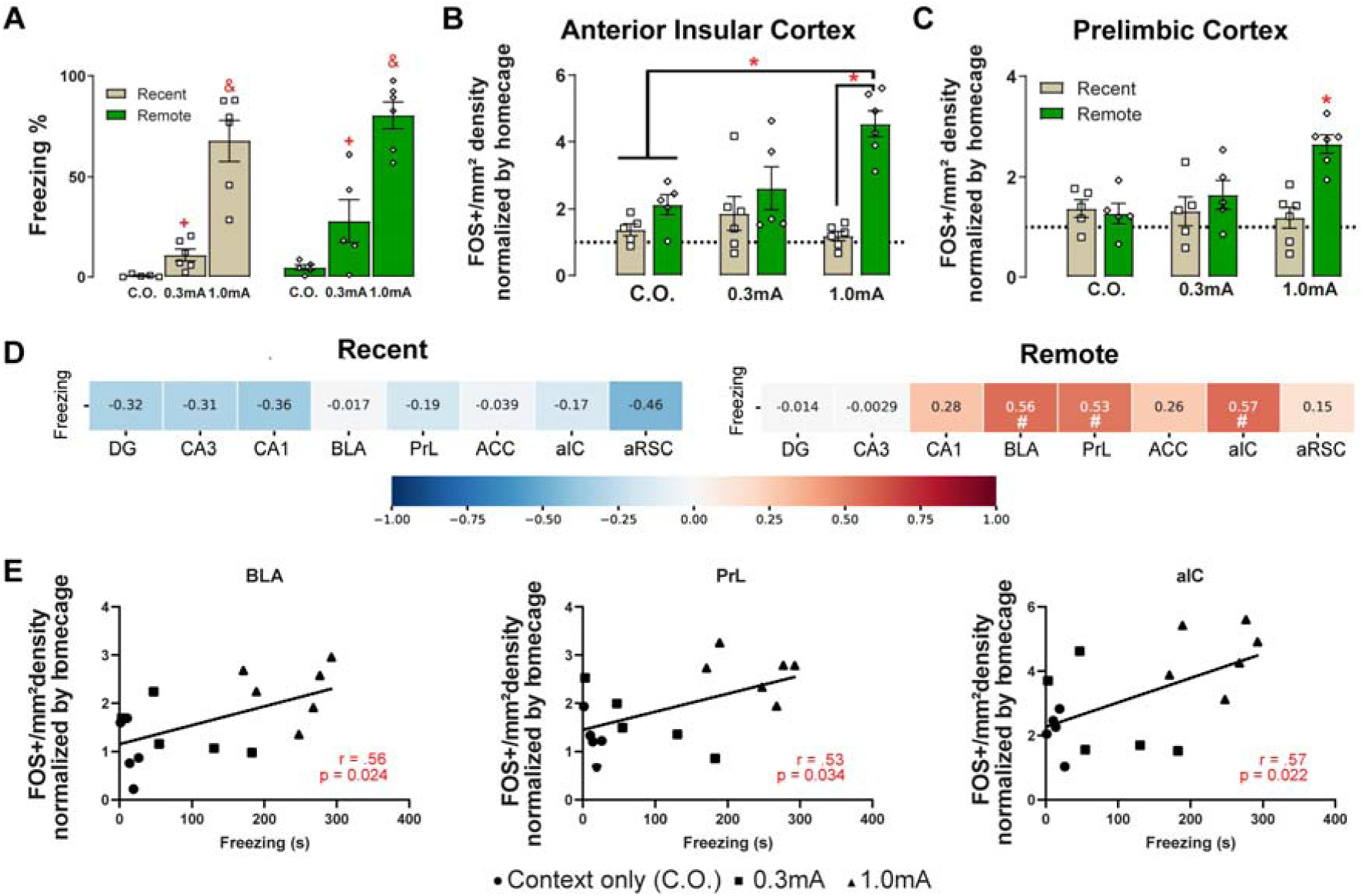
Remote strong contextual fear conditioning retrieval is associated with increased Fos activity in the Anterior Insular and Pre limbic cortices. (a) Mean percentage of freezing time ± Standard Error of the Mean (S.E.M) for rats submitted to the mild (0.3 mA footshocks) or strong (1.0 mA) contextual fear conditioning (CFC) training and tested at the recent and remote timepoints (2 or 28 days after training, respectively). Control animals were exposed to the context but did not receive any footshocks during training and were tested at the same timepoints (Context only, C.O.). Points represent raw data for each rat. N=5 for the 0.3mA|Remote and C.O. groups, N=6 for all other groups. (+) p < 0.05 when compared to the C.O. group at the same timepoint. (&) p < 0.05 when compared to the 0.3mA and C.O. groups at the same timepoint. (b, c) Mean Normalized Fos density ± S.E.M. in the aIC and PrL, respectively. (*) p < 0.05. Symbols show each animal’s raw data. (d) Color-coded heatmap for Pearson correlations between freezing times and Fos activation in all ROIs during recent or remote retrieval. Data from all groups were merged for these tests. Red color represents strong, positive correlations, whereas blue color represents negative ones. (#) significant bicaudal Pearson correlation, p < 0.05, uncorrected. (e) Correlations between freezing times during remote test sessions and Fos activation in the BLA, PrL and aIC. All groups were analyzed together and the Pearson correlation results (r and p-values) are shown in red; triangles correspond to the 1.0mA|Remote group, squares to the 0.3mA|Remote group, and circles to the C.O.|Remote group.

Among the 8 brain regions analyzed, two-way ANOVAs revealed interaction effects for the aIC, PrL and aRSC. Figure 1 (b, c) shows quantification of Fos cells in the aIC and PrL, respectively. In the aIC we observed an increase of Fos signal only in the 1.0 mA|Remote group. The ANOVA showed significant effects for the Timepoint factor: F(1,27) = 10.55, *p* < 0.01, *η*^*2*^*p* = 0.28 and Footshock * Timepoint interaction: F(2,27) = 5.84, *p* < 0.01, *η*^*2*^*p* = 0.30, but not for the Footshock factor: F(2, 27) = 2.40, *p* = 0.11. Post hoc tests showed that the group of rats trained with 1.0 mA intensity and tested on the remote timepoint had a significant increase of immunoreactive cells when compared to rats trained with the same intensity and tested at the recent timepoint (*p* < 0.001, d = 2.75 95%CI = [1.34-4.16]) and to rats only exposed to the context during training and tested at the recent (*p* = 0.003, d = 2.59 95%CI = [1.15-4.03]) or remote (*p* = 0.032, d = 1.98 95%CI = [0.62-3.34]) timepoints. All other comparisons were not significant (*p* > 0.09, Fig. 2b). A similar increase was seen in the PrL. The ANOVA showed significant effects for the Footshock factor: F(2, 26) = 3.89, *p* = 0.033, *η*^*2*^*p* = 0.23; Timepoint factor: F(1,26) = 9.22, *p* < 0.01, *η*^*2*^*p* = 0.26 and Footshock * Timepoint interaction: F(2,26) = 6.58, *p* < 0.01, *η*^*2*^*p* = 0.34. Post hoc tests confirmed that the group of rats trained with 1.0 mA intensity and tested on the remote timepoint had a significant increase of immunoreactive cells when compared to all other groups (*p* < 0.042) whereas all other pairwise comparisons were not significant (*p* > 0.70, Fig. 2c). Table S3 and S4 report all post hoc comparisons for the aIC and PrL ANOVAs, with their effect sizes and 95%CI. Even though the aRSC ANOVA showed an interaction effect [F(2,26) = 3.77, *p* = 0.04, *η*^*2*^*p* = 0.23, Table 2], post hoc tests did not confirm any differences between groups (*p* > 0.10, Table S5). The Footshock and Timepoint factors for the aRSC ANOVA were also not significant (*p* > 0.29, Table 2). Among the other 5 ROIs, two-way ANOVAs revealed main effects of Timepoint for the DG and CA1, but no main effect of Footshock or interactions (see Table 2 for full statistics). The ANOVAs also did not show any significant effects for the BLA, CA3 and ACC (Table 2).

**Table 2.** Descriptive (mean normalized Fos density ± S.E.M) and inferential statistics (ANOVAs) for other analyzed brain regions.

| Brain Region |  | Context-only |  | 0.3mA |  | 1.0mA |  | ANOVA |  |  |  |
| --- | --- | --- | --- | --- | --- | --- | --- | --- | --- | --- | --- |
|  |  | Recent | Remote | Recent | Remote | Recent | Remote | Factor | F | df | p-value |
| **DG** | Mean | 1.50 | 1.94 | 1.31 | 1.74 | 0.877 | 2.22 | Footshock | 0.119 | 2,27 | 0.888 |
| | S.E.M ± | 0.415 | 0.549 | 0.515 | 0.528 | 0.204 | 0.280 | Timepoint | 4.485 | 1,27 | **0.044\*** |
|  | N | 5 | 5 | 6 | 5 | 6 | 6 | Footshock*Timepoint | 0.796 | 2,27 | 0.461 |
| **CA3** | Mean | 1.44 | 1.29 | 1.12 | 0.948 | 0.922 | 1.48 | Footshock | 0.326 | 2,27 | 0.725 |
| | S.E.M ± | 0.459 | 0.553 | 0.388 | 0.509 | 0.310 | 0.172 | Timepoint | 0.062 | 1,27 | 0.805 |
|  | N | 5 | 5 | 6 | 5 | 6 | 6 | Footshock*Timepoint | 0.552 | 2,27 | 0.582 |
| **CA1** | Mean | 1.74 | 2.23 | 1.38 | 1.78 | 0.889 | 3.63 | Footshock | 0.501 | 2,27 | 0.612 |
| | S.E.M ± | 0.528 | 0.886 | 0.508 | 0.904 | 0.217 | 0.915 | Timepoint | 4.517 | 1,27 | **0.043\*** |
|  | N | 5 | 5 | 6 | 5 | 6 | 6 | Footshock*Timepoint | 1.898 | 2,27 | 0.169 |
| **BLA** | Mean | 1.11 | 1.03 | 1.13 | 1.43 | 1.11 | 2.29 | Footshock | 2.27 | 2,27 | 0.123 |
| | S.E.M ± | 0.407 | 0.276 | 0.153 | 0.238 | 0.424 | 0.238 | Timepoint | 3.48 | 1,27 | 0.073 |
|  | N | 5 | 5 | 6 | 5 | 6 | 6 | Footshock*Timepoint | 2.30 | 2,27 | 0.120 |
| **aRSC** | Mean | 1.52 | 0.971 | 1.38 | 1.65 | 1.00 | 2.03 | Footshock | 0.557 | 2,26 | 0.580 |
| | S.E.M ± | 0.175 | 0.111 | 0.285 | 0.437 | 0.194 | 0.310 | Timepoint | 1.151 | 1,26 | 0.293 |
|  | N | 5 | 4 | 6 | 5 | 6 | 6 | Footshock*Timepoint | 3.772 | 2,26 | **0.036\*** |
| **ACC** | Mean | 0.893 | 0.837 | 0.703 | 0.836 | 0.846 | 1.71 | Footshock | 2.21 | 2,27 | 0.129 |
| | S.E.M ± | 0.155 | 0.233 | 0.200 | 0.275 | 0.243 | 0.370 | Timepoint | 2.12 | 1,27 | 0.157 |
|  | N | 5 | 5 | 6 | 5 | 6 | 6 | Footshock*Timepoint | 1.76 | 2,27 | 0.192 |

Note: (\*) refers to a significant effect.

**Figure 2:**
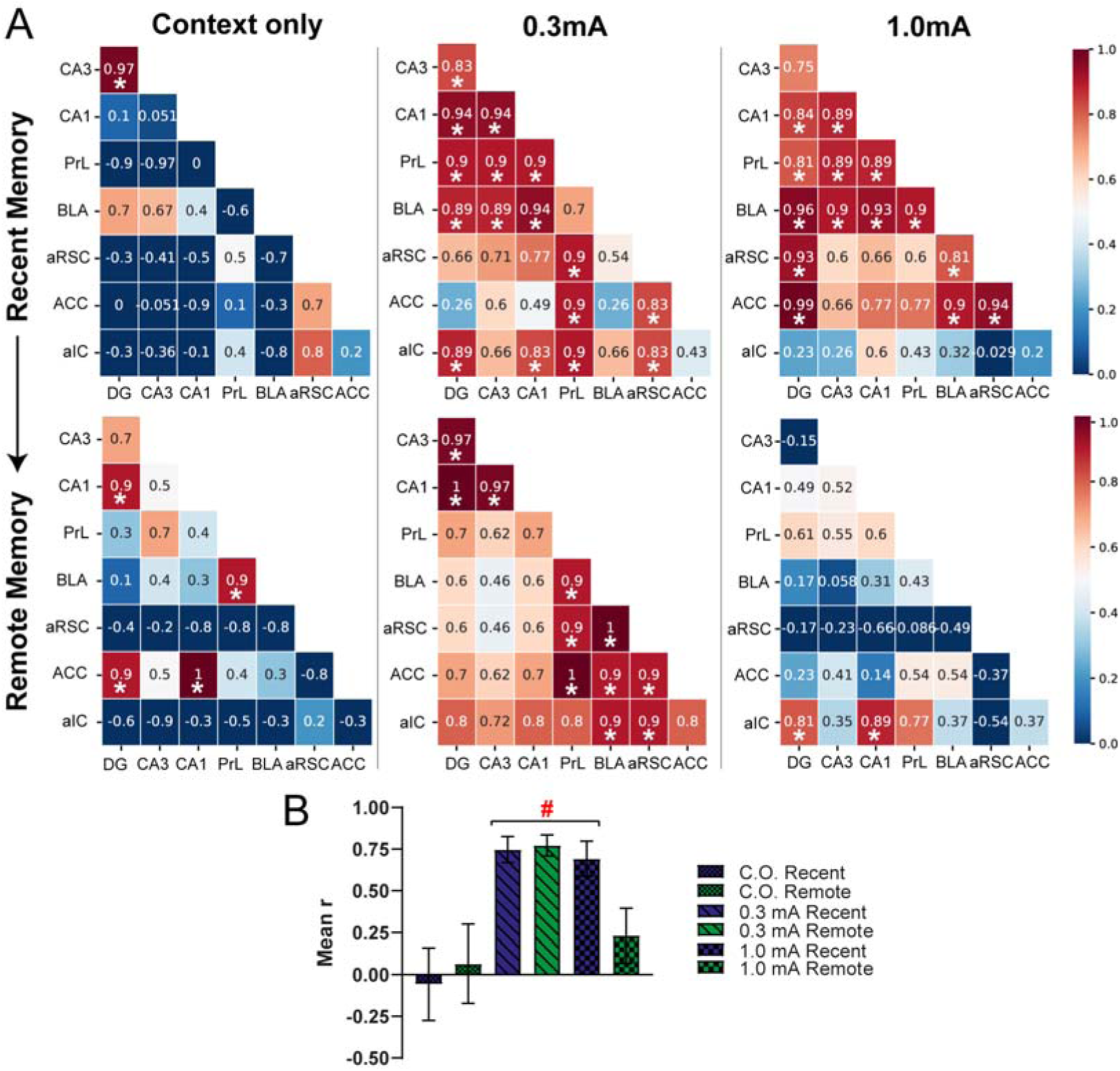
Increasing footshock intensity elicits different trends of functional connectivity between contextual memory brain regions during remote retrieval. (a) Correlation matrices showing pairwise co-variance between Regions of Interest (ROIs), or functional connectivity. Rats were submitted to the mild (0.3 mA footshocks) or strong (1.0 mA) contextual fear conditioning (CFC) training and tested at the recent and remote timepoints (2 or 28 days after training, respectively). Control animals were exposed to the context but did not receive any footshocks during training and were tested at the same timepoints (Context only, C.O.). Each correlation test compared vectors sized 5-6 (number of animals). Red color represents strong positive correlations, whereas blue color represents strong negative correlations (*) p ≤ 0.025 in the unicaudal Spearman correlation test, uncorrected for multiple comparisons. (b) Mean r values ± 95% confidence interval across all ROIs per group. (#) p < 0.001 in comparison to the 1.0mA|Remote and Context Only groups (recent and remote timepoints).

To verify which regions were most active while freezing behavior was expressed, we tested the association between freezing times and Fos activity for each ROI with 16 bicaudal Pearson correlation tests for the recent and remote timepoints (8 per timepoint). All groups exposed to the context were combined for the analyses. The correlation tests did not reveal any significant correlation for the recent timepoint (*p* > 0.07, see Fig. 1d) but revealed significant positive correlations for the BLA, PrL and aIC during remote retrieval (*p* = 0.024, 0.034 and 0.022, respectively, see Fig. 1d, e). All other correlation tests for the remote timepoint were not significant (*p* > 0.43).

In summary, regional immunoreactivity analyses showed an increase in Fos expression during the remote retrieval test in the aIC and PrL for the strong CFC. On the other hand, there was an overall increase in Fos cells for all groups during the remote CFC test in the DG and CA1 hippocampal subregions, regardless of footshock intensity. The correlation tests also suggest that Fos activity in the aIC, PrL and BLA during remote retrieval seems to be positively associated with higher freezing levels.

### 2. Pairwise co-variance between ROIs

To map neuronal coactivity between brain areas implicated in either recent or remote memory retrieval, Fos expression was analyzed in the ROIs from each animal, and interregional Spearman correlations were calculated for each group (split by footshock intensity and timepoint, Figure 2a). This analysis allows identifying pairs of brain regions in which normalized Fos density co-varies between subjects. Only those with high correlation were considered functionally connected (significant correlations rho ≥ 0.81, *p* ≤ 0.025 uncorrected). Results showed that Context Only groups show little to no coactivation between the ROIs in both timepoints. On the other hand, for the conditioned groups tested in the recent timepoint, many ROIs showed coactivation between themselves, in both footshock intensities. It is of note that the BLA and PrL show significant coactivation with all hippocampal subregions at this timepoint. Additionally, during remote retrieval, the 0.3mA group shows sustained high coactivation between ROIs, similar to the pattern shown by the recent group, but with weakened coactivation between the BLA, aIC and PrL with hippocampal subregions. In contrast, the 1.0mA|Remote group showed lessened coactivation between ROIs, having significant results only between the aIC and the DG-CA1.

To determine if coactivation between areas differed between experimental groups, we compared the averaged “r” values of all 8 ROIs using a Welch 1-way ANOVA (Worley et al., 2020). The ANOVA showed a significant effect of Group [F(5,73.3) = 23.2, *p* < 0.001, *η*^2^p = 0.42 Fig. 2b]. Averaged across all ROIs, we found more correlated activity in the 0.3mA|Recent, 0.3mA|Remote, 1.0mA|Recent when compared to the C.O.|Recent, C.O.|Remote and 1.0mA|Remote groups (Fig. 2b; for all post hoc results and effect sizes, see Table S6). To further investigate specific differences in coactivation between the shocked groups from the recent to the remote timepoint, data from each correlation was contrasted between timepoints using the Fisher r to z transformation and z test (Table 3). The test for the 0.3mA groups indicated a significant increase in coactivation over time between the BLA-aRSC and from the DG-CA1. On the other hand, there was a significant decrease in connectivity in the 1.0mA groups between the DG-ACC, BLA-aRSC as well as between the aRSC-ACC and aRSC-BLA (Table 3). P-values of all z tests contrasts can be seen in Table S7. Lastly, we contrasted data from each footshock intensity by timepoint to investigate possible differences in coactivation between the shocked groups (Table 3). For the Recent timepoint, results indicated higher connectivity between the DG and ACC in the 1.0mA compared to the 0.3mA group. Conversely, results from the Remote timepoint indicated that the 0.3mA group had higher connectivity between the DG-CA1 and DG-CA3, as well as between the aRSC-BLA, aRSC-aIC and ACC-PrL. Statistical results for these z tests can be seen in Table S8.

**Table 3.**
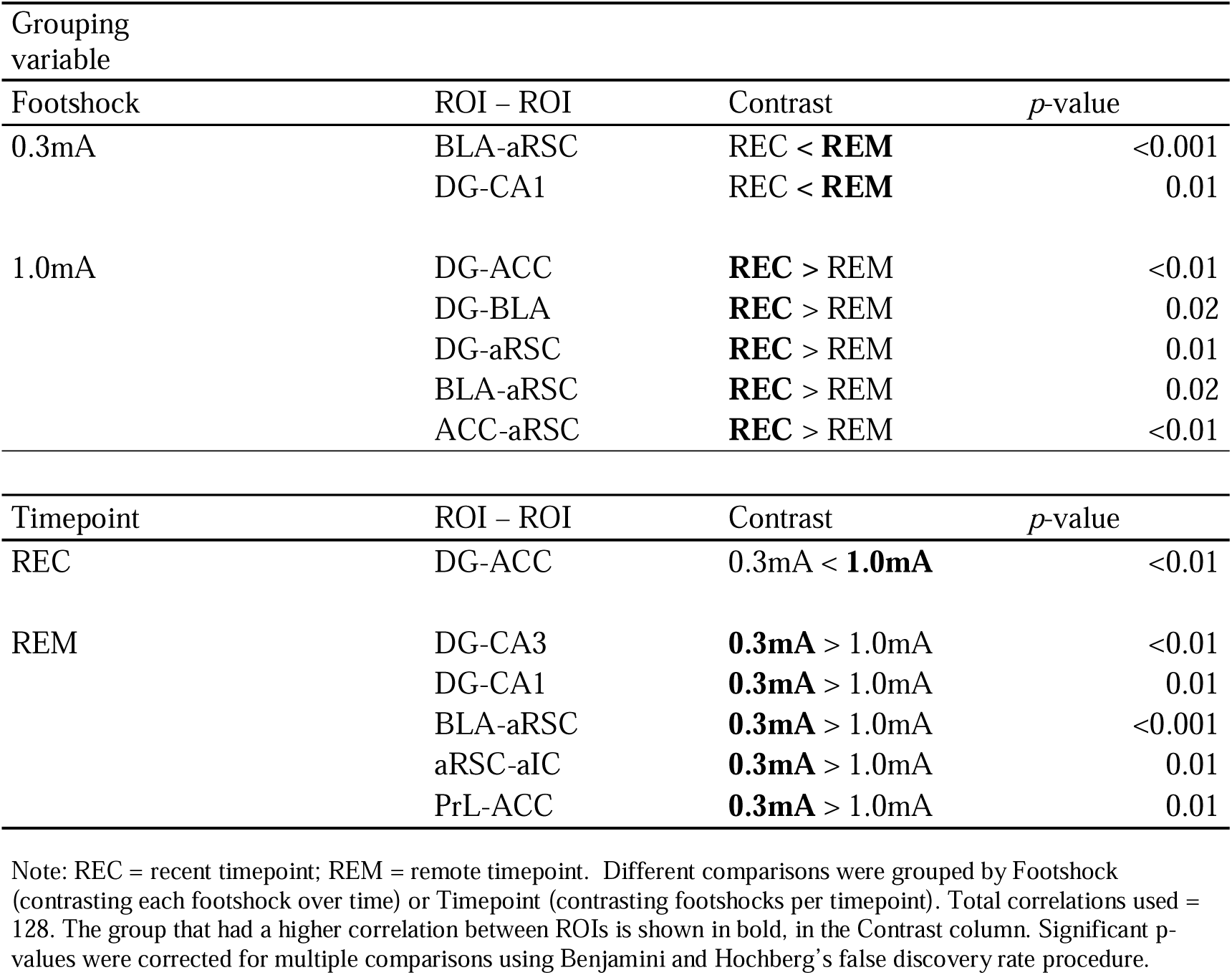
Summary of significant pairwise ROI by ROI comparisons.

| Grouping variable | ROI – ROI | Contrast | *p*-value |
| --- | --- | --- | --- |
| Footshock |  |  |  |
| 0.3mA | BLA-aRSC | REC < **REM** | <0.001 |
|  | DG-CA1 | REC < **REM** | 0.01 |
| 1.0mA | DG-ACC | **REC** > REM | <0.01 |
|  | DG-BLA | **REC** > REM | 0.02 |
|  | DG-aRSC | **REC** > REM | 0.01 |
|  | BLA-aRSC | **REC** > REM | 0.02 |
|  | ACC-aRSC | **REC** > REM | <0.01 |
| Timepoint | ROI – ROI | Contrast | *p*-value |
| REC | DG-ACC | 0.3mA < **1.0mA** | <0.01 |
| REM | DG-CA3 | **0.3mA** > 1.0mA | <0.01 |
|  | DG-CA1 | **0.3mA** > 1.0mA | 0.01 |
|  | BLA-aRSC | **0.3mA** > 1.0mA | <0.001 |
|  | aRSC-aIC | **0.3mA** > 1.0mA | 0.01 |
|  | PrL-ACC | **0.3mA** > 1.0mA | 0.01 |

Note: REC = recent timepoint; REM = remote timepoint. Different comparisons were grouped by Footshock (contrasting each footshock over time) or Timepoint (contrasting footshocks per timepoint). Total correlations used = 128. The group that had a higher correlation between ROIs is shown in bold, in the Contrast column. Significant *p*-values were corrected for multiple comparisons using Benjamini and Hochberg's false discovery rate procedure.

In summary, these results show that increasing footshock intensity in the CFC task elicited different trends in Fos coactivation between cortical and subcortical regions associated with contextual fear memories during remote retrieval, with a significant decrease of overall connectivity between ROIs in the 1.0mA|Remote group compared to all other conditioned groups. Results from this group also show stronger decreases in coactivation between the aRSC and DG to the BLA and ACC when comparing the two timepoints. On the other hand, the 0.3mA footshock intensity seems to maintain high connectivity between ROIs over time. Likewise, the 0.3mA group had higher coactivation between the aRSC to the BLA and aIC, the DG to other hippocampal subareas, and the ACC to the PrL when compared to the 1.0mA at the Remote timepoint.

### 3. Association between post-training CORT levels and Fos expression during recent and remote memory retrieval

Results from the Spearman correlation tests showed that Context Only groups had little to no association between the post-training CORT levels and Fos expression in the recent timepoint (*p* > 0.35, uncorrected. Figure 3a). On the other hand, for the remote timepoint, there were significant positive correlations between CORT and Fos in the CA1 (rho = .99, *p* = 0.02, uncorrected) and ACC (rho= .99, *p* = 0.02, uncorrected). There were no significant correlations for the rats trained with 0.3mA footshocks and tested in both timepoints (*p* > 0.23, uncorrected). On the other hand, the 1.0mA|Recent group shows strong positive correlations between CORT and Fos in the CA3 (rho = .94, *p* = 0.02, uncorrected), CA1 (rho = .94, *p* = 0.02, uncorrected) and BLA (rho = 0.84, *p* = 0.04, uncorrected). A close-to-significance correlation was also found between CORT and Fos in the PrL (rho = .83, *p* = 0.058, uncorrected). In contrast, the 1.0mA|Remote group did not have any significant correlations (*p* > 0.08, uncorrected).

**Figure 3:**
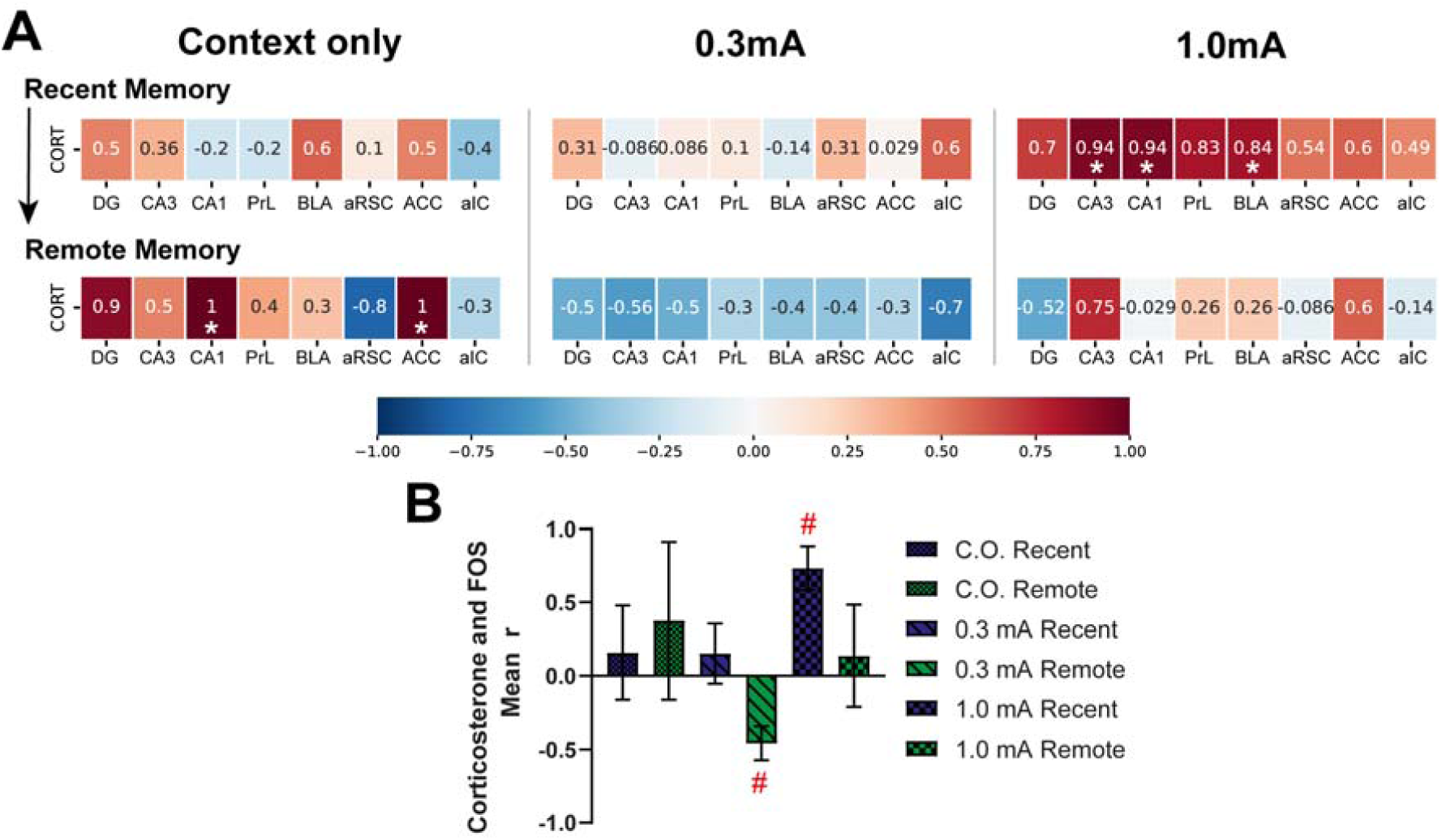
Post-training corticosterone (CORT) levels are differentially associated with neuronal activity during contextual memory retrieval after mild or strong CFC training. (a) Color-coded heatmap for Spearman correlations between post-training CORT levels and Fos activation in all ROIs during recent or remote retrieval, split by timepoint and training intensity. Rats were submitted to the mild (0.3 mA footshocks) or strong (1.0 mA) contextual fear conditioning (CFC) training, had their blood collected 30 minutes after training, and were tested at the recent and remote timepoints (2 or 28 days after training, respectively). Control animals were exposed to the context and blood collection but did not receive any footshocks during training and were tested at the same timepoints (Context only, C.O.). Each correlation test compared vectors sized 5-6 (number of animals). Red color represents strong positive correlations, whereas blue color represents strong negative correlations (*) p < 0.05 in the bicaudal Spearman correlation test, uncorrected for multiple comparisons. (b) Mean r values ± 95% confidence interval across all ROIs per group. (#) p < 0.05 in comparison to all other groups except the C.O. Remote group.

Considering that CORT acts on memory consolidation modulating neuronal activity in several brain regions simultaneously, we investigated whether average correlations differed between experimental groups. We performed a similar analysis to the one described previously using the averaged “r” values of all 8 CORT-Fos correlations with a Welch 1-way ANOVA, that showed a significative effect [F(5,18.9) = 40.6, *p* < 0.001, η^2^p = 0.50, Fig. 3b]. Averaged across all ROIs, we found a moderate inverse relationship between CORT and Fos expression in the 0.3mA|Remote group when compared to the C.O.|Recent, 0.3mA|Recent, 1.0mA|Recent, and 1.0mA|Remote groups (*p* ≤ 0.036). On the other hand, the 1.0mA|Recent group had a strong, positive association with Fos when compared to all other groups (*p* ≤ 0.036) except the C.O.|Remote group (Fig. 3b, for all post hoc results and effect sizes, see Table S9).

In summary, these results show that post-training CORT levels are differentially associated with Fos expression during contextual memory retrieval after mild or strong CFC training. CORT seems to be inversely associated with neuronal activity in the remote retrieval of mild CFC, whereas after strong CFC it showed strong positive correlations with CA3, CA1 and BLA during the recent retrieval.

## DISCUSSION

Our laboratory recently showed that stronger CFC trainings (3x 0.6 or 1.0mA footshocks) elicited time-dependent fear generalization, whereas mild CFC trainings (3x 0.3mA footshocks) prompted remote contextual discrimination (Dos Santos Corrêa et al., 2019). We suggested that these results were due to different systems consolidation processes that occur after a mild or strong CFC, which is now further confirmed here. We also observed that higher CORT levels were associated with time-dependent fear generalization (Dos Santos Corrêa et al., 2019). Now, we demonstrate that strong CFC training elicits higher Fos activation of the aIC and PrL during remote retrieval (Fig. 1b, c), and Fos activity in these regions and in the BLA at this timepoint is associated with freezing response to the training context (Fig. 1d, e). Also, remote retrieval after mild CFC training seems to induce sustained high functional connectivity, whereas strong CFC training elicits lower functional connectivity between the ROIs at this timepoint (Fig. 2). Moreover, connectivity across regions is mostly equivalent between footshocks at the recent timepoint, whereas, at the remote one, there were more substantial differences between mild and strong CFC – especially between the DG and other hippocampal subregions, the aRSC to the BLA and aIC, and the PrL and ACC – implicating in the differential engagement of memory-related brain networks over time according to arousal levels during training (Table 3; Footshock). Likewise, strong CFC remote retrieval seems to have diminished connectivity between the aRSC and DG to the BLA and ACC compared to the same training intensity in recent retrieval, suggesting a possible decrease in engagement of the retrosplenial/hippocampus network overtime after stronger trainings (Table 3; Timepoint).

In addition, although plasma CORT levels after training were similar in rats trained with 1.0 or 0.3 mA footshocks, only those trained with the strongest CFC protocol showed higher CORT levels than the C.O. group (Fig. S2). Furthermore, post-training CORT levels were differentially associated with neuronal activation according to training intensity, indicating that the glucocorticoid might have a modulatory effect on engram allocation during memory consolidation, thus influencing the process of systems consolidation over time (Fig. 3). Interestingly, for the strong CFC protocol, post-training CORT is positively correlated with Fos density in two hippocampal subregions (CA1 and CA3) and in the BLA during recent retrieval (with the PrL showing a trend towards significance, Fig 3a). These regions (hippocampal subregions, the BLA, and PrL) are among the ones that show significant differences in connectivity over time, and between training protocols at the remote timepoint. The overall data suggest that increasing training intensity during memory acquisition results in differential processes of systems consolidation that can be further associated with either fear discrimination after mild CFC or generalized fear after strong CFC, possibly correlated to circulating glucocorticoid levels during memory consolidation (Dos Santos Corrêa et al., 2019). Fos levels reported here are similar to those conveyed in past studies that have analyzed neuronal activity after recent and remote contextual fear memory retrieval (Silva et al., 2019; Tayler et al., 2013).

Wheeler and colleagues (2013) showed that there is increased co-variance between neocortical and hippocampal regions after remote retrieval, in contrast to what is found after recent retrieval when co-variance seems to be restricted to a hippocampal-centered network (Wheeler et al., 2013). Earlier lesion studies indicated that remote retrieval becomes independent of the hippocampus, and the memory trace seems to lose some of the contextual details that are necessary for discriminative retrieval in this process (Frankland and Bontempi, 2005). Nonetheless, this hypothesis has been challenged by several studies showing that the hippocampus remains engaged during remote memory retrieval (Albo and Gräff, 2018; Sutherland and Lehmann, 2011; Vetere et al., 2021). Our results strengthen this last view. Our data suggest that some neocortical regions become more engaged during memory retrieval at remote timepoints (but only for strong CFC) and the hippocampus may maintain its functional connectivity to other ROIs (after mild CFC) at this same timepoint. It is possible that, in the earlier lesion studies, the strengthening of neocortical functional connectivity over time compensated for the absence of a functional hippocampus during remote retrieval, and may be the reason why temporarily inactivating the hippocampus before remote retrieval tests did not impair freezing levels (Quillfeldt, 2019).

Our results suggest an association between freezing behavior and Fos activity in the PrL, BLA and aIC during remote CFC retrieval (Fig. 1c), possibly indicating that engram cells allocated in these regions during the acquisition and consolidation of strong CFC may mature over time into a “contextual fear network”. Interestingly, the BLA and aIC are associated with the SN in rodents (Mandino et al., 2021; Zerbi et al., 2019). Recent evidence has shown that changes in the SN connectivity due to acute stress may be a potential risk factor for the later development of trauma-related symptoms (Zhang et al., 2022). Even though there is some divergence over which region of the mPFC, specifically the ACC or PrL, is indeed part of the SN in rodents (Gozzi and Schwarz, 2016; Hermans et al., 2014b; Seeley, 2019), recent studies have reported that the PrL shows functional connectivity to both the SN and DMN-like (Mandino et al., 2021; Tsai et al., 2020) possibly having differential engagement to these networks in more or less aversive events. On the other hand, the aRSC and ACC (again, including both the ACC and PrL, but especially the ACC) in rodents are usually associated with the DMN-like network (Gozzi and Schwarz, 2016; Grandjean et al., 2020; Mandino et al., 2021), an analogous of the DMN in humans, which is inversely correlated with SN activation (Zerbi et al., 2015) especially after acute stress (Upadhyay et al., 2011). Accordingly, our z-scores results (Table 3) demonstrate that the main differences in connectivity across the Footshock and Timepoint factors happen between the aRSC and ACC to the BLA, PrL and hippocampal subregions, indicating that these regions may become less functionally connected in stronger CFC or remote retrieval. Therefore, we hypothesize that strong contextual fear memory may engage some brain regions associated with the SN during the consolidation phase, especially the aIC and PrL, via increased noradrenergic and glucocorticoid activity in these regions (Schwabe et al., 2022), and possibly this strong CFC engram inhibits the engagement of DMN-like regions such as the ACC and aRSC over time. This hypothesis is strengthened by evidence from a recent elegant study that verified an inhibitory pathway from the CA3 to the anterodorsal thalamic nucleus (ADn), which matures over time and is necessary for remote memory retrieval (Vetere et al., 2021). This thalamic nucleus seems to be part of the DMN-like network (Grandjean et al., 2020; Mandino et al., 2021), receives projections from the ACC, aRSC and presubiculum (Vetere et al., 2021), and the circuitry between these regions and its effects on memory are already well described (Bubb et al., 2017; Mitchell et al., 2018). It is possible that this inhibitory pathway from the CA3 to the ADn is differentially engaged during mild or strong CFC, facilitating the disengagement of the DMN-like observed in the 1.0mA|Remote group via the ADn, but other experiments would be necessary to confirm this supposition.

Other evidence from neuroimaging studies in humans have associated increased coherence between the aIC, mPFC and amygdala in response to stressful events or treatment with noradrenergic or glucocorticoid agonists (Quaedflieg and Schwabe, 2018). Correspondingly, recent studies with PTSD patients have indicated that they show an exacerbated processing of interoceptive/arousal stimuli in the aIC (Nicholson et al., 2020), increased connectivity between the ACC and the aIC (Caseras et al., 2013) and also that these brain regions are associated with generalized threat detection in these patients (Berg et al., 2020). Webler and colleagues (2021) described that an increasing gradient of fear memory generalization is associated with higher connectivity between the aIC, dorsomedial PFC (the analogous of the rodent PrL), dorsal ACC along with other regions, and also with a decrease in connectivity in nodes of the DMN as generalization increased (Webler et al., 2021). Therefore, it is possible that strongly arousing events (i.e. strong CFC training) produce an enduring memory engram in neocortical regions associated with the SN, which are initially silent and mature over time (Josselyn and Tonegawa, 2020), undergoing systems consolidation. On the other hand, a different process seems to occur after mildly arousing events, maintaining memory dependence on DMN-like regions.

Although there seems to be a consensus that a memory trace undergoes systems consolidation, it has been debated which brain regions are initially recruited to allocate engram cells in different memory tasks (Chaaya et al., 2018). Moreover, it is unclear when memory retrieval would become independent of hippocampal subregions and more dependent on neocortical ones (Frankland and Bontempi, 2005). Recent evidence demonstrates that a brain-wide engram circuit is allocated after memory acquisition (Rao-Ruiz et al., 2021; Roy et al., 2022). Previous studies showed that for the CFC task there is engram allocation in the hippocampus, mPFC (including the prelimbic cortex and the ACC), aRSC and amygdala (Matos et al., 2019; Ohkawa et al., 2015). Our data suggest that engram cells may also be allocated in the aIC during memory acquisition, but the engagement of this region over time could be especially dependent on training intensity, possibly due to modulatory influence from glucocorticoid activity (Fornari et al., 2012). Additionally, a recent study has also shown that CORT treatment immediately after fear conditioning increases engram size in DG during memory consolidation, and chemogenetically inactivating this engram during recent retrieval elicits lower generalized fear in a novel context (Lesuis et al., 2021), indicating that, at least for this region, the glucocorticoid amplifies engram allocation. The same modulatory role can be true for other regions, such as the mPFC and the amygdala. Even though the study on the relationship between post-training CORT levels and engram allocation is still incipient, there are several studies indicating a modulatory role played by the glucocorticoid system on memory strength (Roozendaal, 2002) and specificity (Bahtiyar et al., 2020; Gazarini et al., 2021). Our results suggest that circulating CORT levels during consolidation may act on several brain regions simultaneously modulating not only engram allocation but also the trajectory of systems consolidation, albeit our data is only correlational and more studies are needed to confirm possible causal effects. Data presented here can also explain the differential effects seen in contextual specificity with post-training CORT treatments after mild or moderate CFC trainings, in which CORT seems to promote contextual discrimination in mild CFC up to 28 days and to facilitate fear generalization after moderate CFC at 14 days (Dos Santos Corrêa et al., 2021), and explain the contextual generalization effects seen in the inhibitory avoidance task (Roozendaal and Mirone, 2020).

Besides the possible modulatory effect glucocorticoids have in engram allocation and systems consolidation, they seem to also interact with arousal levels and the noradrenergic system. Recent studies in rodents showed that noradrenergic activation of the BLA during memory consolidation influences neural plasticity in the aIC and PrL and enhances recognition memory (Barsegyan et al., 2019). Likewise, activation of glucocorticoid receptors in the aIC is associated with enhanced contextual fear memory consolidation (Fornari et al., 2012). These results suggest that aversive events may recruit these regions during memory acquisition and consolidation. However, is this recruitment arousal-dependent? It seems likely to be. The engagement of the aIC in the retrieval of CFC is intensity-dependent, as its post-training functional inhibition impairs recent retrieval after stronger trainings (Alves et al., 2013) but not weaker ones (de Paiva et al., 2021). This dependence on the training intensity was also reported in the PrL in mice (Matos et al., 2019). These results together agree with ours and suggest that the aIC and PrL may be recruited differentially for the consolidation of contextual memories after mild or stronger aversive events. Lastly, the increase in functional connectivity between DG and CA1 in the 1.0mA|Remote group to the aIC is intriguing, considering that there are few input and output projections to and from the aIC and the hippocampus (Gehrlach et al., 2020). Possibly, functional connectivity between these two regions may be explained by their mutual projections to the medial prefrontal cortex, suggesting that a mPFC-aIC pathway is engaged during strong CFC remote retrieval. Barsegyan and colleagues (2019) provide evidence that the PrL may serve as a site of integration between the hippocampal and insular systems (Barsegyan et al., 2019) and this circuit might also be relevant for strong remote CFC retrieval.

A limitation of our study is that we have only quantified Fos from animals that were tested in the training context and not in a novel one. More experiments are needed to investigate neuronal activation in the ROIs that are engaged during exposure to novel contexts after mild or strong CFC. It is possible that the fear network investigated here is engaged only when the animal is exposed to the training context, therefore all associations between generalized fear and loss of functional connectivity between the ROIs studied here are second-order inferences. Moreover, the lack of differences between rats trained with or without footshocks for most ROIs is noteworthy. Even though we expected to see differences in the activation of hippocampal and amygdalar regions, this difference was not confirmed by the data. Another study that also quantified Fos expression has found a lack of difference between these groups in the dorsal hippocampus at the remote timepoint (Silva et al., 2019) but this study and others have found significant differences in neocortical and amygdalar regions that we did not, especially in the ACC and aRSC (Aceti et al., 2015; Silva et al., 2019). These studies, however, used several different parameters that could account for the divergent results, such as the use of a different species (mice), stronger training parameters (5 or 30 footshocks), or a different behavioral task (inhibitory avoidance). Another limitation is the small sample size per experimental group which decreases statistical power for correlations tests, but this limitation was mitigated by choosing a unicaudal correlation test, uncorrected for multiple comparisons due to a large number of repeated tests (Ben-Ami Bartal et al., 2021).

To conclude, we found correlational evidence between an increase of Fos activation in regions associated with the SN after strong CFC remote retrieval. Hence, we infer that increasing arousal levels (with contiguous increased glucocorticoid signaling in the brain) during memory consolidation elicit different processes of systems consolidation, resulting in the recruitment of different clusters of brain regions in response to contextual fear conditioning. Strong CFC elicits the release of higher levels of CORT (Dos Santos Corrêa et al., 2019), and maybe noradrenaline, possibly recruiting regions associated with the SN (PrL and aIC). On the other hand, mild CFC training would not reach the necessary levels of arousal that allow engram allocation in these areas, preserving contextually-specific remote retrieval (Dos Santos Corrêa et al., 2021; Pedraza et al., 2016), and the aforementioned dependence on the DMN-like. The highest footshock intensity training was previously associated with time-dependent fear generalization, implicating a possible relationship between increased neuronal activity in the SN to generalized freezing (Dos Santos Corrêa et al., 2019). Overgeneralization of fear is one of the most debilitating symptoms of PTSD and generalized anxiety disorder, consequently our results indicate the necessity of targeting the SN in future neuromodulatory treatments for these disorders.

## Acknowledgments

We thank the valuable support from Bruna de Oliveira, Thaisy Moraes Costa and Gabriel Leir Gandra for helping in some experiments. We also thank the work done by the technicians that manage the vivarium. The authors would like to thank the members of the Research group in Neurobiology of Learning and Memory at UFABC (MANAs) for scientific advice and discussion.

## Research data for this article

The data used for all analyses in this study can be found at https://data.mendeley.com/datasets/kztbft87y6.

## Author Statement

Moisés dos Santos Corrêa: Conceptualization, Methodology, Data curation, Formal analysis, Investigation, Resources, Visualization, Writing – original draft. Gabriel David Vieira Grisanti: Validation, Investigation, Writing-Reviewing and Editing. Isabelle Anjos Fernandes Franciscatto: Validation, Investigation, Writing-Reviewing and Editing. Tatiana Suemi Anglas Tarumoto: Validation, Investigation, Writing-Reviewing and Editing. Paula Ayako Tiba: Conceptualization, Funding acquisition, Methodology, Supervision, Writing – review & editing. Tatiana Lima Ferreira: Conceptualization, Funding acquisition, Methodology, Supervision, Investigation, Writing – review & editing. Raquel Vecchio Fornari: Conceptualization, Funding acquisition, Methodology, Supervision, Investigation, Writing – review & editing, Project administration.

## REFERENCES

[dataset] dos Santos Corrêa, M., Barbara dos Santos, V., Gabriel David Vieira, G., Joselisa Péres Queiroz, de P., Paula Ayako, T., Fornari, R.V., 2019. Relationship between footshock intensity, post-training corticosterone release and contextual fear memory specificity over time (raw data). Mendeley Data, v5. https://doi.org/10.17632/n5zdf72x6s.5

Aceti, M., Vetere, G., Novembre, G., Restivo, L., Ammassari-Teule, M., 2015. Progression of activity and structural changes in the anterior cingulate cortex during remote memory formation. Neurobiol. Learn. Mem. 123, 67–71. https://doi.org/10.1016/j.nlm.2015.05.003

Albo, Z., Gräff, J., 2018. The mysteries of remote memory. Philos. Trans. R. Soc. Lond. B. Biol. Sci. 373, 20170029. https://doi.org/10.1098/rstb.2017.0029

Alves, F.H.F., Gomes, F. V., Reis, D.G., Crestani, C.C., Corr, F.M.A., Corrêa, F.M.A., Guimarães, F.S., Resstel, L.B.M., 2013. Involvement of the insular cortex in the consolidation and expression of contextual fear conditioning. Eur. J. Neurosci. 38, 2300–2307. https://doi.org/10.1111/ejn.12210

Arnsten, A.F.T., 2009. Stress signalling pathways that impair prefrontal cortex structure and function. Nat. Rev. Neurosci. 10, 410–422. https://doi.org/10.1038/nrn2648

Atucha, E., Vukojevic, V., Fornari, R. V., Ronzoni, G., Demougin, P., Peter, F., Atsak, P., Coolen, M.W., Papassotiropoulos, A., McGaugh, J.L., de Quervain, D.J.-F., Roozendaal, B., 2017. Noradrenergic activation of the basolateral amygdala maintains hippocampus-dependent accuracy of remote memory. Proc. Natl. Acad. Sci. 201710819. https://doi.org/10.1073/pnas.1710819114

Bahtiyar, S., Karaca, K.G., Henckens, M.J.A.G.A.G., Roozendaal, B., Gulmez Karaca, K., Henckens, M.J.A.G.A.G., Roozendaal, B., 2020. Norepinephrine and glucocorticoid effects on the brain mechanisms underlying memory accuracy and generalization. Mol. Cell. Neurosci. 108, 103537. https://doi.org/10.1016/j.mcn.2020.103537

Barry, D.N., Maguire, E.A., 2019. Remote Memory and the Hippocampus: A Constructive Critique. Trends Cogn. Sci. 23, 128–142. https://doi.org/10.1016/j.tics.2018.11.005

Barsegyan, A., Mirone, G., Ronzoni, G., Guo, C., Song, Q., van Kuppeveld, D., Schut, E.H.S., Atsak, P., Teurlings, S., McGaugh, J.L., Schubert, D., Roozendaal, B., 2019. Glucocorticoid enhancement of recognition memory via basolateral amygdala-driven facilitation of prelimbic cortex interactions. Proc. Natl. Acad. Sci. U. S. A. 116, 7077–7082. https://doi.org/10.1073/pnas.1901513116

Ben-Ami Bartal, I., Breton, J.M., Sheng, H., Long, K.L.P.L., Chen, S., Halliday, A., Kenney, J.W., Wheeler, A.L., Frankland, P., Shilyansky, C., Deisseroth, K., Keltner, D., Kaufer, D., Bartal, I.B.A., Breton, J.M., Sheng, H., Long, K.L.P.L., Chen, S., Halliday, A., Kenney, J.W., Wheeler, A.L., Frankland, P., Shilyansky, C., Deisseroth, K., Keltner, D., Kaufer, D., 2021. Neural correlates of ingroup bias for prosociality in rats. Elife 10, 1–26. https://doi.org/10.7554/eLife.65582

Berg, H., Ma, Y., Rueter, A., Kaczkurkin, A., Burton, P.C., Deyoung, C.G., Macdonald, A.W., Sponheim, S.R., Lissek, S.M., 2020. Salience and central executive networks track overgeneralization of conditioned-fear in post-traumatic stress disorder. Psychol. Med. https://doi.org/10.1017/S0033291720001166

Bubb, E.J., Kinnavane, L., Aggleton, J.P., 2017. Hippocampal–diencephalic–cingulate networks for memory and emotion: An anatomical guide. Brain Neurosci. Adv. 1, 239821281772344. https://doi.org/10.1177/2398212817723443

Careaga, M.B.L., Girardi, C.E.N., Suchecki, D., 2019. Variability in response to severe stress: highly reactive rats exhibit changes in fear and anxiety-like behavior related to distinct neuronal co-activation patterns. Behav. Brain Res. 373, 112078. https://doi.org/10.1016/j.bbr.2019.112078

Caseras, X., Murphy, K., Mataix-Cols, D., López-Solà, M., Soriano-Mas, C., Ortriz, H., Pujol, J., Torrubia, R., 2013. Anatomical and functional overlap within the insula and anterior cingulate cortex during interoception and phobic symptom provocation. Hum. Brain Mapp. 34, 1220–9. https://doi.org/10.1002/hbm.21503

Chaaya, N., Battle, A.R., Johnson, L.R., 2018. An update on contextual fear memory mechanisms: Transition between Amygdala and Hippocampus. Neurosci. Biobehav. Rev. 92, 43–54. https://doi.org/10.1016/j.neubiorev.2018.05.013

Coelho, C.A.O., Ferreira, T.L., Kramer-Soares, J.C., Sato, J.R., Oliveira, M.G.M., 2018. Network supporting contextual fear learning after dorsal hippocampal damage has increased dependence on retrosplenial cortex. PLoS Comput. Biol. 14, e1006207. https://doi.org/10.1371/journal.pcbi.1006207

de Paiva, J.P.Q., Bueno, A.P.A., Dos Santos Corrêa, M., Oliveira, M.G.M., Ferreira, T.L., Fornari, R. V, 2021. The posterior insular cortex is necessary for the consolidation of tone fear conditioning. Neurobiol. Learn. Mem. 179, 107402. https://doi.org/10.1016/j.nlm.2021.107402

DeNardo, L.A., Liu, C.D., Allen, W.E., Adams, E.L., Friedmann, D., Fu, L., Guenthner, C.J., Tessier-Lavigne, M., Luo, L., 2019. Temporal evolution of cortical ensembles promoting remote memory retrieval. Nat. Neurosci. 22, 460–469. https://doi.org/10.1038/s41593-018-0318-7

Dos Santos Corrêa, M., Vaz, B. dos S., Grisanti, G.D.V., de Paiva, J.P.Q., Tiba, P.A., Fornari, R.V., 2019. Relationship between footshock intensity, post-training corticosterone release and contextual fear memory specificity over time. Psychoneuroendocrinology 110, 104447. https://doi.org/10.1016/j.psyneuen.2019.104447

Dos Santos Corrêa, M., Vaz, B. dos S., Menezes, B.S., Ferreira, T.L., Tiba, P.A., Fornari, R.V., 2021. Corticosterone differentially modulates time-dependent fear generalization following mild or moderate fear conditioning training in rats. Neurobiol. Learn. Mem. 184, 107487. https://doi.org/10.1016/j.nlm.2021.107487

Fornari, R. V., Wichmann, R., Atucha, E., Desprez, T., Eggens-Meijer, E., Roozendaal, B., 2012. Involvement of the insular cortex in regulating glucocorticoid effects on memory consolidation of inhibitory avoidance training. Front. Behav. Neurosci. 6, 10. https://doi.org/10.3389/fnbeh.2012.00010

Frankland, P.W., Bontempi, B., 2005. The organization of recent and remote memories. Nat. Rev. Neurosci. 6, 119–30. https://doi.org/10.1038/nrn1607

Frankland, P.W., Bontempi, B., Talton, L.E., Kaczmarek, L., Silva, A.J., 2004. The Involvement of the Anterior Cingulate Cortex in Remote Contextual Fear Memory. Science (80-.). 304, 881–883. https://doi.org/10.1126/science.1094804

Gazarini, L., Stern, C.A., Takahashi, R.N., Bertoglio, L.J., 2021. Interactions of Noradrenergic, Glucocorticoid and Endocannabinoid Systems Intensify and Generalize Fear Memory Traces. Neuroscience. https://doi.org/10.1016/j.neuroscience.2021.09.012

Gehrlach, D.A., Weiand, C., Gaitanos, T.N., Cho, E., Klein, A.S., Hennrich, A.A., Conzelmann, K., Gogolla, N., 2020. A whole-brain connectivity map of mouse insular cortex. Elife 9, 2020.02.10.941518. https://doi.org/10.7554/eLife.55585

Gozzi, A., Schwarz, A.J., 2016. Large-scale functional connectivity networks in the rodent brain. Neuroimage 127, 496–509. https://doi.org/10.1016/j.neuroimage.2015.12.017

Grandjean, J., Canella, C., Anckaerts, C., AyrancI, G., Bougacha, S., Bienert, T., Buehlmann, D., Coletta, L., Gallino, D., Gass, N., Garin, C.M., Nadkarni, N.A., Hübner, N.S., Karatas, M., Komaki, Y., Kreitz, S., Mandino, F., Mechling, A.E., Sato, C., Sauer, K., Shah, D., Strobelt, S., Takata, N., Wank, I., Wu, T., Yahata, N., Yeow, L.Y., Yee, Y., Aoki, I., Chakravarty, M.M., Chang, W.T., Dhenain, M., von Elverfeldt, D., Harsan, L.A., Hess, A., Jiang, T., Keliris, G.A., Lerch, J.P., Meyer-Lindenberg, A., Okano, H., Rudin, M., Sartorius, A., Van der Linden, A., Verhoye, M., Weber-Fahr, W., Wenderoth, N., Zerbi, V., Gozzi, A., 2020. Common functional networks in the mouse brain revealed by multi-centre resting-state fMRI analysis. Neuroimage 205. https://doi.org/10.1016/j.neuroimage.2019.116278

Hermans, E.J., Battaglia, F.P., Atsak, P., De Voogd, L.D., Fernández, G., Roozendaal, B., 2014a. How the amygdala affects emotional memory by altering brain network properties. Neurobiol. Learn. Mem. 112, 2–16. https://doi.org/10.1016/j.nlm.2014.02.005

Hermans, E.J., Henckens, M.J.A.G.A.G., Joëls, M., Fernández, G., 2014b. Dynamic adaptation of large-scale brain networks in response to acute stressors. Trends Neurosci. 37, 304–314. https://doi.org/10.1016/j.tins.2014.03.006

Ji, M., Xia, J., Tang, X., Yang, J., 2018. Altered functional connectivity within the default mode network in two animal models with opposing episodic memories. PLoS One 13, e0202661. https://doi.org/10.1371/journal.pone.0202661

Josselyn, S.A., Tonegawa, S., 2020. Memory engrams: Recalling the past and imagining the future. Science 367. https://doi.org/10.1126/science.aaw4325

Lesuis, S.L., Brosens, N., Immerzeel, N., van der Loo, R.J., Mitrić, M., Bielefeld, P., Fitzsimons, C.P., Lucassen, P.J., Kushner, S.A., van den Oever, M.C., Krugers, H.J., 2021. Glucocorticoids promote fear generalization by increasing the size of a dentate gyrus engram cell population. Biol. Psychiatry 1446, 2669. https://doi.org/10.1016/j.biopsych.2021.04.010

Mahan, A.L., Ressler, K.J., 2012. Fear conditioning, synaptic plasticity and the amygdala: Implications for posttraumatic stress disorder. Trends Neurosci. 35, 24–35. https://doi.org/10.1016/j.tins.2011.06.007

Mandino, F., Vrooman, R.M., Foo, H.E., Yeow, L.Y., Bolton, T.A.W., Salvan, P., Teoh, C.L., Lee, C.Y., Beauchamp, A., Luo, S., Bi, R., Zhang, J., Lim, G.H.T., Low, N., Sallet, J., Gigg, J., Lerch, J.P., Mars, R.B., Olivo, M., Fu, Y., Grandjean, J., 2021. A triple-network organization for the mouse brain. Mol. Psychiatry 1–8. https://doi.org/10.1038/s41380-021-01298-5

Matos, M.R., Visser, E., Kramvis, I., van der Loo, R.J., Gebuis, T., Zalm, R., Rao-Ruiz, P., Mansvelder, H.D., Smit, A.B., van den Oever, M.C., 2019. Memory strength gates the involvement of a CREB-dependent cortical fear engram in remote memory. Nat. Commun. 10, 2315. https://doi.org/10.1038/s41467-019-10266-1

Mitchell, A.S., Czajkowski, R., Zhang, N., Jeffery, K., Nelson, A.J.D., 2018. Retrosplenial cortex and its role in spatial cognition. Brain Neurosci. Adv. 2, 239821281875709. https://doi.org/10.1177/2398212818757098

Moscarello, J.M., Maren, S., 2018. Flexibility in the face of fear: hippocampal–prefrontal regulation of fear and avoidance. Curr. Opin. Behav. Sci. 19, 44–49. https://doi.org/10.1016/j.cobeha.2017.09.010

Moscovitch, M., Cabeza, R., Winocur, G., Nadel, L., 2016. Episodic memory and beyond: The hippocampus and neocortex in transformation. Annu. Rev. Psychol. 67, 105–134. https://doi.org/10.1146/annurev-psych-113011-143733

Nicholson, A.A., Harricharan, S., Densmore, M., Neufeld, R.W.J., Ros, T., McKinnon, M.C., Frewen, P.A., Théberge, J., Jetly, R., Pedlar, D., Lanius, R.A., 2020. Classifying heterogeneous presentations of PTSD via the default mode, central executive, and salience networks with machine learning. NeuroImage Clin. 27, 102262. https://doi.org/10.1016/j.nicl.2020.102262

Ohkawa, N., Saitoh, Y., Suzuki, A., Tsujimura, S., Murayama, E., Kosugi, S., Nishizono, H., Matsuo, M., Takahashi, Y., Nagase, M., Sugimura, Y.K., Watabe, A.M., Kato, F., Inokuchi, K., 2015. Artificial association of pre-stored information to generate a qualitatively new memory. Cell Rep. 11, 261–269. https://doi.org/10.1016/j.celrep.2015.03.017

Paxinos, G., Watson, C.R., 2007. The Rat Brain in Stereotaxic Coordinates, 6th Editio. ed. Academic Press.

Pedraza, L.K., Sierra, R.O., Boos, F.Z., Haubrich, J., Quillfeldt, J.A., de Oliveira Alvares, L., 2016. The dynamic nature of systems consolidation: Stress during learning as a switch guiding the rate of the hippocampal dependency and memory quality. Hippocampus 26, 362–371. https://doi.org/10.1002/hipo.22527

Quaedflieg, C.W.E.M., Schwabe, L., 2018. Memory dynamics under stress. Memory 26, 364–376. https://doi.org/10.1080/09658211.2017.1338299

Quillfeldt, J.A., 2019. Temporal Flexibility of Systems Consolidation and the Synaptic Occupancy/Reset Theory (SORT): Cues About the Nature of the Engram. Front. Synaptic Neurosci. 11, 1. https://doi.org/10.3389/fnsyn.2019.00001

Rao-Ruiz, P., Visser, E., Mitrić, M., Smit, A.B., van den Oever, M.C., 2021. A Synaptic Framework for the Persistence of Memory Engrams. Front. Synaptic Neurosci. 13, 1–15. https://doi.org/10.3389/fnsyn.2021.661476

Redondo, R.L., Kim, J., Arons, A.L., Ramirez, S., Liu, X., Tonegawa, S., 2014. Bidirectional switch of the valence associated with a hippocampal contextual memory engram. Nature 513, 426–430. https://doi.org/10.1038/nature13725

Roozendaal, B., 2002. Stress and Memory: Opposing Effects of Glucocorticoids on Memory Consolidation and Memory Retrieval GLUCOCORTICOIDS AND MEMORY FUNCTION. Neurobiol. Learn. Mem. 78, 578–595. https://doi.org/10.1006/nlme.2002.4080

Roozendaal, B., Mirone, G., 2020. Opposite Effects of Noradrenergic and Glucocorticoid Activation on Accuracy of Episodic-like Memory. Psychoneuroendocrinology 114, 104588. https://doi.org/10.1016/j.psyneuen.2020.104588

Roy, D.S., Arons, A., Mitchell, T.I., Pignatelli, M., Ryan, T.J., Tonegawa, S., 2016. Memory retrieval by activating engram cells in mouse models of early Alzheimer’s disease. Nature 531, 508–512. https://doi.org/10.1038/nature17172

Roy, D.S., Park, Y.G., Kim, M.E., Zhang, Y., Ogawa, S.K., DiNapoli, N., Gu, X., Cho, J.H., Choi, H., Kamentsky, L., Martin, J., Mosto, O., Aida, T., Chung, K., Tonegawa, S., 2022. Brain-wide mapping reveals that engrams for a single memory are distributed across multiple brain regions. Nat. Commun. 13. https://doi.org/10.1038/s41467-022-29384-4

Sano, Y., Shobe, J.L., Zhou, M., Huang, S., Shuman, T., Cai, D.J., Golshani, P., Kamata, M., Silva, A.J., 2014. CREB regulates memory allocation in the insular cortex. Curr. Biol. 24, 2833–2837. https://doi.org/10.1016/j.cub.2014.10.018

Schwabe, L., 2017. Memory under stress: from single systems to network changes. Eur. J. Neurosci. 45, 478–489. https://doi.org/10.1111/ejn.13478

Schwabe, L., Hermans, E.J., Joëls, M., Roozendaal, B., 2022. Mechanisms of memory under stress. Neuron 1–18. https://doi.org/10.1016/j.neuron.2022.02.020

Seeley, W.W., 2019. The salience network: A neural system for perceiving and responding to homeostatic demands. J. Neurosci. 39, 9878–9882. https://doi.org/10.1523/JNEUROSCI.1138-17.2019

Silva, B.A., Burns, A.M., Gräff, J., 2019. A cFos activation map of remote fear memory attenuation. Psychopharmacology (Berl). 236, 369–381. https://doi.org/10.1007/s00213-018-5000-y

Sutherland, R.J., Lehmann, H., 2011. Alternative conceptions of memory consolidation and the role of the hippocampus at the systems level in rodents. Curr. Opin. Neurobiol. 21, 446–451. https://doi.org/10.1016/j.conb.2011.04.007

Tayler, K.K., Tanaka, K.Z., Reijmers, L.G., Wiltgen, B.J., 2013. Reactivation of neural ensembles during the retrieval of recent and remote memory. Curr. Biol. 23, 99–106. https://doi.org/10.1016/j.cub.2012.11.019

Tsai, P.J., Keeley, R.J., Carmack, S.A., Vendruscolo, J.C.M., Lu, H., Gu, H., Vendruscolo, L.F., Koob, G.F., Lin, C.P., Stein, E.A., Yang, Y., 2020. Converging Structural and Functional Evidence for a Rat Salience Network. Biol. Psychiatry 88, 867–878. https://doi.org/10.1016/j.biopsych.2020.06.023

Upadhyay, J., Baker, S.J., Chandran, P., Miller, L., Lee, Y., Marek, G.J., Sakoglu, U., Chin, C.L., Luo, F., Fox, G.B., Day, M., 2011. Default-Mode-Like Network Activation in Awake Rodents. PLoS One 6. https://doi.org/10.1371/journal.pone.0027839

Vetere, G., Xia, F., Ramsaran, A.I., Tran, L.M., Josselyn, S.A., Frankland, P.W., 2021. An inhibitory hippocampal–thalamic pathway modulates remote memory retrieval. Nat. Neurosci. 24, 685–693. https://doi.org/10.1038/s41593-021-00819-3

Webler, R.D., Berg, H., Fhong, K., Tuominen, L., Holt, D.J., Morey, R.A., Lange, I., Burton, P.C., Fullana, M.A., Radua, J., Lissek, S., 2021. The neurobiology of human fear generalization: meta-analysis and working neural model. Neurosci. Biobehav. Rev. 128, 421–436. https://doi.org/10.1016/j.neubiorev.2021.06.035

Wheeler, A.L., Teixeira, C.M., Wang, A.H., Xiong, X., Kovacevic, N., Lerch, J.P., McIntosh, A.R., Parkinson, J., Frankland, P.W., 2013. Identification of a Functional Connectome for Long-Term Fear Memory in Mice. PLoS Comput. Biol. 9. https://doi.org/10.1371/journal.pcbi.1002853

Winocur, G., Moscovitch, M., Bontempi, B., 2010. Memory formation and long-term retention in humans and animals: Convergence towards a transformation account of hippocampal-neocortical interactions. Neuropsychologia 48, 2339–2356. https://doi.org/10.1016/j.neuropsychologia.2010.04.016

Worley, N.B., Everett, S.R., Foilb, A.R., Christianson, J.P., 2020. Functional networks activated by controllable and uncontrollable stress in male and female rats. Neurobiol. Stress 13, 100233. https://doi.org/10.1016/j.ynstr.2020.100233

Zerbi, V., Floriou-Servou, A., Markicevic, M., Vermeiren, Y., Sturman, O., Privitera, M., von Ziegler, L., Ferrari, K.D., Weber, B., De Deyn, P.P., Wenderoth, N., Bohacek, J., 2019. Rapid Reconfiguration of the Functional Connectome after Chemogenetic Locus Coeruleus Activation. Neuron 103, 702-718.e5. https://doi.org/10.1016/j.neuron.2019.05.034

Zerbi, V., Grandjean, J., Rudin, M., Wenderoth, N., 2015. Mapping the mouse brain with rs-fMRI: An optimized pipeline for functional network identification. Neuroimage 123, 11–21. https://doi.org/10.1016/j.neuroimage.2015.07.090

Zhang, W., Kaldewaij, R., Hashemi, M.M., Koch, S.B.J., Smit, A., van Ast, V.A., Beckmann, C.F., Klumpers, F., Roelofs, K., 2022. Acute-stress-induced change in salience network coupling prospectively predicts post-trauma symptom development. Transl. Psychiatry 12, 1–8. https://doi.org/10.1038/s41398-022-01798-0

Zhang, Y., Fukushima, H., Kida, S., 2011. Induction and requirement of gene expression in the anterior cingulate cortex and medial prefrontal cortex for the consolidation of inhibitory avoidance memory. Mol. Brain 4, 4. https://doi.org/10.1186/1756-6606-4-4

