## Supplemental Material for "Remote contextual fear retrieval engages activity from salience network regions in rats"

**Supplementary Material for “REMOTE CONTEXTUAL FEAR RETRIEVAL ENGAGES ACTIVITY FROM SALIENCE NETWORK REGIONS IN RATS”**

**Table S1– Post hoc test results for freezing behavior in recent and remote memory retrieval tests**

| Post Hoc Comparisons - Recent | | | | | | | | | | |
| --- | --- | --- | --- | --- | --- | --- | --- | --- | --- | --- |
| **Comparison** | | |  | | | | | | **95% Confidence Interval** | |
| **Footshock** |  | **Footshock** | **Mean Difference** | **SE** | **df** | **t** | **p_tukey_** | **Cohen's d** | **Lower** | **Upper** |
| C.O. | - | 0.3ma | -4.46 | 1.30 | 14.0 | -3.43 | 0.011 | -2.08 | -3.62 | -0.529 |
|  | - | 1.0ma | -13.01 | 1.30 | 14.0 | -10.00 | < .001 | -6.06 | -8.84 | -3.280 |
| 0.3ma | - | 1.0ma | -8.55 | 1.24 | 14.0 | -6.90 | < .001 | -3.98 | -6.02 | -1.947 |
| *Note.* Comparisons are based on estimated marginal means | | | | | | | | | | |
| Post Hoc Comparisons - Remote | | | | | | | | | | |
| **Comparison** | | |  | | | | | | **95% Confidence Interval** | |
| **Footshock** |  | **Footshock** | **Mean Difference** | **SE** | **df** | **t** | **p_tukey_** | **Cohen's d** | **Lower** | **Upper** |
| C.O. | - | 0.3ma | -5.17 | 1.71 | 14.0 | -3.01 | 0.024 | -1.82 | -3.32 | -0.330 |
|  | - | 1.0ma | -11.92 | 1.71 | 14.0 | -6.95 | < .001 | -4.21 | -6.35 | -2.064 |
| 0.3ma | - | 1.0ma | -6.75 | 1.64 | 14.0 | -4.13 | 0.003 | -2.38 | -3.95 | -0.812 |
| *Note.* Comparisons are based on estimated marginal means | | | | | | | | | | |

**Table S2 and Figure S1 – One-way ANOVA for Plasma Corticosterone between footshock intensities**

| ANOVA - SQRT Cort | | | | | | |  |  |  |  |
| --- | --- | --- | --- | --- | --- | --- | --- | --- | --- | --- |
|  | **Sum of Squares** | **df** | **Mean Square** | **F** | **p** | **η²p** |  |  |  |  |
| Footshock | 128 | 2 | 63.9 | 3.40 | 0.046 | 0.180 |  |  |  |  |
| Residuals | 584 | 31 | 18.8 |  |  |  |  |  |  |  |
| Post Hoc Comparisons - Footshock | | | | | | | | | | |
| **Comparison** | | |  | | | | | | **95% Confidence Interval** | |
| **Footshock** |  | **Footshock** | **Mean Difference** | **SE** | **df** | **t** | **p_tukey_** | **Cohen's d** | **Lower** | **Upper** |
| C.O. | - | 0.3ma | -3.14 | 1.86 | 31.0 | -1.691 | 0.224 | -0.724 | -1.62 | 0.169 |
|  | - | 1.0ma | -4.80 | 1.86 | 31.0 | -2.584 | 0.038 | -1.107 | -2.03 | -0.188 |
| 0.3ma | - | 1.0ma | -1.66 | 1.77 | 31.0 | -0.937 | 0.622 | -0.382 | -1.22 | 0.456 |
| *Note.* Comparisons are based on estimated marginal means | | | | | | | | | | |

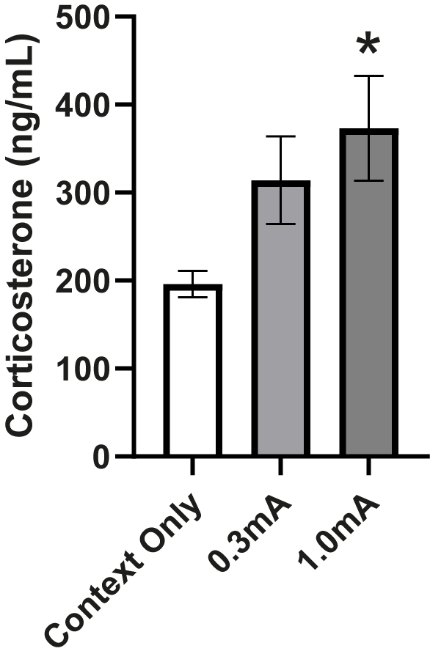
Figure S1: Mean plasma corticosterone (ng/mL) ± S.E.M per experimental group. The subset of rats used in this study is representative of the results published previously (Dos Santos Corrêa et al., 2019). The animals trained with 1.0mA footshocks show increased CORT levels when compared to the context only group. (*) p < 0,05 in comparison to the Context Only group. Number of animals per group: Context Only: 10, 0.3mA: 12, 1.0mA: 12.

Dos Santos Corrêa, M., Vaz, B. dos S., Grisanti, G.D.V., de Paiva, J.P.Q., Tiba, P.A., Fornari, R.V., 2019. Relationship between footshock intensity, post-training corticosterone release and contextual fear memory specificity over time. Psychoneuroendocrinology 110, 104447. https://doi.org/10.1016/j.psyneuen.2019.104447

**Table S3 – Post hoc test results for the anterior Insular Cortex factorial ANOVA**

| Anterior Insular Cortex Post Hoc Comparisons - Footshock ✻ Timepoint | | | | | | | | | | | | |
| --- | --- | --- | --- | --- | --- | --- | --- | --- | --- | --- | --- | --- |
| **Comparison** | | | | |  | | | | | | **95% Confidence Interval** | |
| **Footshock** | **Timepoint** |  | **Footshock** | **Timepoint** | **Mean Difference** | **SE** | **df** | **t** | **p_tukey_** | **Cohen's d** | **Lower** | **Upper** |
| C.O. | REC | - | C.O. | REM | -0.7516 | 0.771 | 27 | -0.9744 | 0.922 | -0.6163 | -1.9253 | 0.6930 |
|  |  | - | 0.3ma | REC | -1.1934 | 0.738 | 27 | -1.6161 | 0.596 | -0.9786 | -2.2507 | 0.2940 |
|  |  | - | 0.3ma | REM | -1.2467 | 0.771 | 27 | -1.6163 | 0.595 | -1.0223 | -2.3510 | 0.3060 |
|  |  | - | 1.0ma | REC | 0.1883 | 0.738 | 27 | 0.2550 | 1.000 | 0.1544 | -1.0888 | 1.3980 |
|  |  | - | 1.0ma | REM | -3.1622 | 0.738 | 27 | -4.2822 | 0.003 | -2.5930 | -4.0310 | -1.1550 |
|  | REM | - | 0.3ma | REC | -0.4418 | 0.738 | 27 | -0.5983 | 0.990 | 0.3623 | -0.8842 | 1.6090 |
|  |  | - | 0.3ma | REM | -0.4951 | 0.771 | 27 | -0.6419 | 0.987 | -0.4060 | -1.7086 | 0.8970 |
|  |  | - | 1.0ma | REC | 0.9399 | 0.738 | 27 | 1.2728 | 0.797 | -0.7707 | -2.0317 | 0.4900 |
|  |  | - | 1.0ma | REM | -2.4107 | 0.738 | 27 | -3.2644 | 0.032 | -1.9767 | -3.3362 | -0.6170 |
| 0.3ma | REC | - | 0.3ma | REM | -0.0532 | 0.738 | 27 | -0.0721 | 1.000 | -0.0437 | -1.2862 | 1.1990 |
|  |  | - | 1.0ma | REC | 1.3817 | 0.704 | 27 | 1.9625 | 0.389 | 1.1330 | -0.0931 | 2.3590 |
|  |  | - | 1.0ma | REM | -1.9688 | 0.704 | 27 | -2.7962 | 0.089 | -1.6144 | -2.8819 | -0.3470 |
|  | REM | - | 1.0ma | REC | 1.4350 | 0.738 | 27 | 1.9432 | 0.399 | -1.1767 | -2.4618 | 0.1080 |
|  |  | - | 1.0ma | REM | -1.9156 | 0.738 | 27 | -2.5940 | 0.133 | -1.5708 | -2.8883 | -0.2530 |
| 1.0ma | REC | - | 1.0ma | REM | -3.3506 | 0.704 | 27 | -4.7587 | < .001 | -2.7474 | -4.1588 | -1.3360 |
| *Note.* Comparisons are based on estimated marginal means | | | | | | | | | | | | |

**Table S4 – Post hoc test results for the Prelimbic Cortex factorial ANOVA**

| Pre Limbic Post Hoc Comparisons - Footshock ✻ Timepoint | | | | | | | | | | | | |
| --- | --- | --- | --- | --- | --- | --- | --- | --- | --- | --- | --- | --- |
| **Comparison** | | | | |  | | | | | | **95% Confidence Interval** | |
| **Footshock** | **Timepoint** |  | **Footshock** | **Timepoint** | **Mean Difference** | **SE** | **df** | **t** | **p_tukey_** | **Cohen's d** | **Lower** | **Upper** |
| C.O. | REC | - | C.O. | REM | 0.0974 | 0.330 | 26 | 0.295 | 1 | 0.1866 | -1.115 | 1.488 |
|  |  | - | 0.3ma | REC | 0.0517 | 0.330 | 26 | 0.157 | 1 | 0.0991 | -1.201 | 1.399 |
|  |  | - | 0.3ma | REM | -0.2788 | 0.330 | 26 | -0.845 | 0.956 | -0.5342 | -1.843 | 0.775 |
|  |  | - | 1.0ma | REC | 0.1797 | 0.316 | 26 | 0.569 | 0.992 | 0.3443 | -0.904 | 1.593 |
|  |  | - | 1.0ma | REM | -1.2740 | 0.316 | 26 | -4.032 | 0.005 | -2.4413 | -3.867 | -1.015 |
|  | REM | - | 0.3ma | REC | -0.0457 | 0.330 | 26 | -0.138 | 1 | 0.0875 | -1.213 | 1.388 |
|  |  | - | 0.3ma | REM | -0.3761 | 0.330 | 26 | -1.140 | 0.860 | -0.7207 | -2.037 | 0.595 |
|  |  | - | 1.0ma | REC | 0.0823 | 0.316 | 26 | 0.261 | 1 | -0.1578 | -1.403 | 1.088 |
|  |  | - | 1.0ma | REM | -1.3714 | 0.316 | 26 | -4.340 | 0.002 | -2.6278 | -4.081 | -1.175 |
| 0.3ma | REC | - | 0.3ma | REM | -0.3304 | 0.330 | 26 | -1.001 | 0.913 | -0.6332 | -1.946 | 0.679 |
|  |  | - | 1.0ma | REC | 0.1280 | 0.316 | 26 | 0.405 | 0.998 | 0.2453 | -1.001 | 1.492 |
|  |  | - | 1.0ma | REM | -1.3257 | 0.316 | 26 | -4.195 | 0.003 | -2.5403 | -3.980 | -1.100 |
|  | REM | - | 1.0ma | REC | 0.4584 | 0.316 | 26 | 1.451 | 0.697 | -0.8785 | -2.148 | 0.391 |
|  |  | - | 1.0ma | REM | -0.9952 | 0.316 | 26 | -3.149 | 0.042 | -1.9071 | -3.265 | -0.549 |
| 1.0ma | REC | - | 1.0ma | REM | -1.4537 | 0.301 | 26 | -4.825 | < .001 | -2.7856 | -4.213 | -1.358 |
| *Note.* Comparisons are based on estimated marginal means | | | | | | | | | | | | |

**Table S5 – Post hoc test results for the Anterior Retrosplenial Cortex factorial ANOVA**

| Anterior Retrosplenial Cortex Post Hoc Comparisons - Footshock ✻ Timepoint | | | | | | | | | | | | |
| --- | --- | --- | --- | --- | --- | --- | --- | --- | --- | --- | --- | --- |
| **Comparison** | | | | |  | | | | | | **95% Confidence Interval** | |
| **Footshock** | **Timepoint** |  | **Footshock** | **Timepoint** | **Mean Difference** | **SE** | **df** | **t** | **p_tukey_** | **Cohen's d** | **Lower** | **Upper** |
| C.O. | REC | - | C.O. | REM | 0.5514 | 0.437 | 26 | 1.2625 | 0.802 | 0.8469 | -0.553 | 2.247 |
|  |  | - | 0.3ma | REC | 0.1409 | 0.394 | 26 | 0.3574 | 0.999 | 0.2164 | -1.030 | 1.463 |
|  |  | - | 0.3ma | REM | -0.1312 | 0.412 | 26 | -0.3187 | 1.000 | -0.2016 | -1.503 | 1.100 |
|  |  | - | 1.0ma | REC | 0.5179 | 0.394 | 26 | 1.3136 | 0.775 | 0.7954 | -0.470 | 2.061 |
|  |  | - | 1.0ma | REM | -0.5104 | 0.394 | 26 | -1.2946 | 0.785 | -0.7839 | -2.048 | 0.481 |
|  | REM | - | 0.3ma | REC | -0.4105 | 0.420 | 26 | -0.9768 | 0.921 | 0.6305 | -0.708 | 1.969 |
|  |  | - | 0.3ma | REM | -0.6827 | 0.437 | 26 | -1.5630 | 0.629 | -1.0485 | -2.459 | 0.362 |
|  |  | - | 1.0ma | REC | -0.0335 | 0.420 | 26 | -0.0798 | 1.000 | 0.0515 | -1.275 | 1.378 |
|  |  | - | 1.0ma | REM | -1.0618 | 0.420 | 26 | -2.5265 | 0.153 | -1.6308 | -3.037 | -0.225 |
| 0.3ma | REC | - | 0.3ma | REM | -0.2721 | 0.394 | 26 | -0.6902 | 0.981 | -0.4180 | -1.668 | 0.832 |
|  |  | - | 1.0ma | REC | 0.3770 | 0.376 | 26 | 1.0029 | 0.913 | 0.5790 | -0.619 | 1.777 |
|  |  | - | 1.0ma | REM | -0.6513 | 0.376 | 26 | -1.7326 | 0.524 | -1.0003 | -2.221 | 0.220 |
|  | REM | - | 1.0ma | REC | 0.6491 | 0.394 | 26 | 1.6465 | 0.577 | -0.9970 | -2.274 | 0.280 |
|  |  | - | 1.0ma | REM | -0.3791 | 0.394 | 26 | -0.9617 | 0.926 | -0.5823 | -1.838 | 0.673 |
| 1.0ma | REC | - | 1.0ma | REM | -1.0283 | 0.376 | 26 | -2.7355 | 0.102 | -1.5793 | -2.849 | -0.310 |
| *Note.* Comparisons are based on estimated marginal means | | | | | | | | | | | | |

**Table S6 – Post hoc test results for the averaged “r” values of all 8 ROIs using a Welch 1-way ANOVA**

| Games-Howell Post-Hoc Test – Average Correlations of ROIs per group | | | | | | | | | | | | | |
| --- | --- | --- | --- | --- | --- | --- | --- | --- | --- | --- | --- | --- | --- |
|  | |  | **0.3mARecent** | **0.3mARemote** | | **1.0mARecent** | | **1.0mARemote** | | **CORecent** | | **CORemote** | |
| 0.3mARecent |  | Mean difference | — | -0.0248 |  | 0.0555 |  | 0.516 | *** | 0.805 | *** | 0.683 | *** |
|  |  | t-value | — | -0.507 |  | 0.864 |  | 5.83 |  | 7.17 |  | 5.632 |  |
|  |  | df | — | 52.0 |  | 49.4 |  | 38.5 |  | 33.8 |  | 32.7 |  |
|  |  | p-value | — | 0.996 |  | 0.953 |  | < .001 |  | < .001 |  | < .001 |  |
| 0.3mARemote |  | Mean difference |  | — |  | 0.0802 |  | 0.541 | *** | 0.829 | *** | 0.708 | *** |
|  |  | t-value |  | — |  | 1.328 |  | 6.30 |  | 7.53 |  | 5.932 |  |
|  |  | df |  | — |  | 44.0 |  | 34.9 |  | 31.6 |  | 30.9 |  |
|  |  | p-value |  | — |  | 0.768 |  | < .001 |  | < .001 |  | < .001 |  |
| 1.0mARecent |  | Mean difference |  |  |  | — |  | 0.461 | *** | 0.749 | *** | 0.628 | *** |
|  |  | t-value |  |  |  | — |  | 4.83 |  | 6.36 |  | 4.966 |  |
|  |  | df |  |  |  | — |  | 46.3 |  | 39.3 |  | 37.5 |  |
|  |  | p-value |  |  |  | — |  | < .001 |  | < .001 |  | < .001 |  |
| 1.0mARemote |  | Mean difference |  |  |  |  |  | — |  | 0.289 |  | 0.167 |  |
|  |  | t-value |  |  |  |  |  | — |  | 2.18 |  | 1.191 |  |
|  |  | df |  |  |  |  |  | — |  | 50.3 |  | 48.1 |  |
|  |  | p-value |  |  |  |  |  | — |  | 0.266 |  | 0.839 |  |
| CORecent |  | Mean difference |  |  |  |  |  |  |  | — |  | -0.122 |  |
|  |  | t-value |  |  |  |  |  |  |  | — |  | -0.777 |  |
|  |  | df |  |  |  |  |  |  |  | — |  | 53.6 |  |
|  |  | p-value |  |  |  |  |  |  |  | — |  | 0.970 |  |
| CORemote |  | Mean difference |  |  |  |  |  |  |  |  |  | — |  |
|  |  | t-value |  |  |  |  |  |  |  |  |  | — |  |
|  |  | df |  |  |  |  |  |  |  |  |  | — |  |
|  |  | p-value |  |  |  |  |  |  |  |  |  | — |  |
| *Note.* * p < .05. ** p < .01. *** p < .001 | | | | | | | | | | | | | |

**Table S7 – All pairwise ROI by ROI comparisons between footshock intensities by timepoint**

| Summary of all pairwise ROI by ROI per timepoint comparisons. split by footshock | | | | | | | | | |
| --- | --- | --- | --- | --- | --- | --- | --- | --- | --- |
| **0.3mA** | ROI vs ROI | z score Contrast | p-value | BH FDR | **1.0mA** | ROI vs ROI | z score Contrast | p-value | BH FDR |
|  | **BLA-RSC** | **3.5** | **0.0002** | **0.009** |  | **DG-ACC** | **-2.73** | **0.0032** | **0.009** |
|  | **DG-CA1** | **2.23** | **0.0129** | **0.018** |  | **DG-BLA** | **-2.64** | **0.0174** | **0.018** |
|  | BLA-ACC | 1.32 | 0.0934 | 0.027 |  | **DG-aRSC** | **-2.23** | **0.0129** | **0.027** |
|  | PL-ACC | 1.29 | 0.0985 | 0.036 |  | **BLA-aRSC** | **-2.11** | **0.0207** | **0.036** |
|  | CA1-BLA | -1.17 | 0.121 | 0.045 |  | **ACC-aRSC** | **-2.04** | **0.0041** | **0.045** |
|  | CA3-BLA | -0.99 | 0.1611 | 0.054 |  | CA1-RSC | -1.93 | 0.0268 | 0.054 |
|  | DG-CA3 | 0.97 | 0.166 | 0.063 |  | CA3-BLA | -1.72 | 0.0427 | 0.063 |
|  | CA3-PL | -0.83 | 0.2033 | 0.071 |  | CA1-BLA | -1.61 | 0.0537 | 0.071 |
|  | DG-BLA | -0.78 | 0.2177 | 0.080 |  | DG-CA3 | -1.38 | 0.0838 | 0.080 |
|  | BLA-aIC | 0.75 | 0.2266 | 0.089 |  | PL-BLA | -1.23 | 0.1093 | 0.089 |
|  | ACC-aIC | 0.7 | 0.242 | 0.098 |  | CA3-RSC | -1.14 | 0.1271 | 0.098 |
|  | DG-PL | -0.66 | 0.2546 | 0.107 |  | DG-aIC | 1.1 | 0.1357 | 0.107 |
|  | DG-ACC | 0.66 | 0.2546 | 0.116 |  | CA1-ACC | -1.08 | 0.1401 | 0.116 |
|  | CA1-PL | -0.66 | 0.2546 | 0.125 |  | BLA-ACC | -1.05 | 0.1469 | 0.125 |
|  | PL-BLA | 0.66 | 0.2546 | 0.134 |  | CA3-CA1 | -1.01 | 0.1562 | 0.134 |
|  | CA3-CA1 | 0.45 | 0.3264 | 0.143 |  | CA3-PL | -0.96 | 0.1685 | 0.143 |
|  | CA3-RSc | -0.43 | 0.3336 | 0.152 |  | PL-RSC | -0.95 | 0.1711 | 0.152 |
|  | PL-aIC | -0.41 | 0.3409 | 0.161 |  | CA1-PL | -0.87 | 0.1922 | 0.161 |
|  | CA1-ACC | 0.37 | 0.3557 | 0.170 |  | CA1-aIC | 0.87 | 0.1922 | 0.170 |
|  | CA1-RSC | -0.36 | 0.3594 | 0.179 |  | DG-CA1 | -0.84 | 0.2005 | 0.179 |
|  | DG-aIC | -0.33 | 0.3707 | 0.188 |  | RSC-aIC | -0.71 | 0.2389 | 0.188 |
|  | RSC-ACC | 0.32 | 0.3745 | 0.196 |  | PL-aIC | 0.69 | 0.2451 | 0.196 |
|  | RSC-aIC | 0.32 | 0.3745 | 0.205 |  | DG-PL | -0.52 | 0.3015 | 0.205 |
|  | CA3-aIC | 0.13 | 0.4483 | 0.214 |  | PL-ACC | -0.51 | 0.305 | 0.214 |
|  | DG-RSC | -0.1 | 0.4602 | 0.223 |  | CA3-ACC | -0.44 | 0.33 | 0.223 |
|  | CA1-aIC | -0.09 | 0.4641 | 0.232 |  | ACC-aIC | 0.23 | 0.409 | 0.232 |
|  | CA3-ACC | 0.03 | 0.488 | 0.241 |  | CA3-aIC | 0.12 | 0.4522 | 0.241 |
|  | PL-RSC | 0 | 0.5 | 0.250 |  | BLA-aIC | 0.07 | 0.4721 | 0.250 |
| Note: Contrast order was Remote->Recent. unicaudal. P-values in bold are considered significant after the Benjamini and Hochberg’s false discovery rate (BH FDR) procedure | | | | | | | | | |

**Table S8 – All pairwise ROI by ROI comparisons by timepoint per footshock**

| Summary of all pairwise ROI by ROI per footshock comparisons. split by timepoint | | | | | | | | | |
| --- | --- | --- | --- | --- | --- | --- | --- | --- | --- |
| **REC** | ROI vs ROI | z score Contrast | p-value | BH FDR | **REM** | ROI vs ROI | z score Contrast | p-value | BH FDR |
|  | **DG-ACC** | **2.7** | **0.0035** | 0.00893 |  | **BLA-aRSC** | **-3.48** | **0.0003** | 0.00893 |
|  | aRSC-aIC | -1.48 | 0.0694 | 0.01786 |  | **DG-CA3** | **-2.55** | **0.0054** | 0.01786 |
|  | BLA-ACC | 1.47 | 0.0708 | 0.02679 |  | **DG-CA1** | **-2.31** | **0.0104** | 0.02679 |
|  | DG-aIC | -1.43 | 0.0764 | 0.03571 |  | **aRSC-aIC** | **-2.28** | **0.0113** | 0.03571 |
|  | PL-aIC | -1.24 | 0.1075 | 0.04464 |  | **PL-ACC** | **-2.23** | **0.0129** | 0.04464 |
|  | DG-aRSC | 1.05 | 0.1469 | 0.05357 |  | aRSC-ACC | -2.04 | 0.0207 | 0.05357 |
|  | PL-aRSC | -0.95 | 0.1711 | 0.0625 |  | CA3-CA1 | -1.75 | 0.0401 | 0.0625 |
|  | PL-BLA | 0.73 | 0.2327 | 0.07143 |  | PL-aRSC | -1.71 | 0.0436 | 0.07143 |
|  | aRSC-ACC | 0.71 | 0.2389 | 0.08036 |  | CA1-aRSC | -1.62 | 0.0526 | 0.08036 |
|  | DG-CA1 | -0.66 | 0.2546 | 0.08929 |  | BLA-aIC | -1.19 | 0.117 | 0.08929 |
|  | CA3-aIC | -0.64 | 0.2611 | 0.09821 |  | PL-BLA | -1.11 | 0.1335 | 0.09821 |
|  | BLA-aRSC | 0.64 | 0.2611 | 0.10714 |  | DG-aRSC | -0.95 | 0.1711 | 0.10714 |
|  | DG-BLA | 0.61 | 0.2709 | 0.11607 |  | BLA-ACC | -0.95 | 0.1711 | 0.11607 |
|  | CA1-ACC | 0.6 | 0.2743 | 0.125 |  | CA3-aRSC | -0.81 | 0.209 | 0.125 |
|  | CA1-aIC | -0.6 | 0.2743 | 0.13393 |  | CA1-ACC | -0.79 | 0.2148 | 0.13393 |
|  | BLA-aIC | -0.56 | 0.2877 | 0.14286 |  | ACC-aIC | -0.78 | 0.2177 | 0.14286 |
|  | PL-ACC | -0.55 | 0.2912 | 0.15179 |  | DG-ACC | -0.69 | 0.2451 | 0.15179 |
|  | CA3-CA1 | -0.44 | 0.33 | 0.16071 |  | CA3-aIC | -0.59 | 0.2776 | 0.16071 |
|  | DG-PL | -0.42 | 0.3372 | 0.16964 |  | DG-BLA | -0.57 | 0.2843 | 0.16964 |
|  | ACC-aIC | -0.31 | 0.3783 | 0.17857 |  | CA3-BLA | -0.48 | 0.3156 | 0.17857 |
|  | CA1-aRSC | -0.29 | 0.3859 | 0.1875 |  | CA1-BLA | -0.4 | 0.3446 | 0.1875 |
|  | DG-CA3 | -0.25 | 0.4013 | 0.19643 |  | CA1-aIC | 0.33 | 0.3707 | 0.19643 |
|  | CA3-aRSC | -0.25 | 0.4013 | 0.20536 |  | CA3-ACC | -0.31 | 0.3783 | 0.20536 |
|  | CA1-BLA | -0.15 | 0.4404 | 0.21429 |  | CA1-PL | -0.19 | 0.4247 | 0.21429 |
|  | CA3-ACC | 0.12 | 0.4522 | 0.22321 |  | DG-PL | -0.18 | 0.4286 | 0.22321 |
|  | CA3-PL | -0.09 | 0.4641 | 0.23214 |  | CA3-PL | -0.11 | 0.4562 | 0.23214 |
|  | CA1-PL | -0.09 | 0.4641 | 0.24107 |  | PL-aIC | -0.08 | 0.4681 | 0.24107 |
|  | CA3-BLA | 0.08 | 0.4681 | 0.25 |  | DG-aIC | 0.04 | 0.484 | 0.25 |
| Note: REC = Recent timepoint. REM = Remote timepoint. Contrast order was 1.0mA->0.3mA. unicaudal. P-values in bold are considered significant after the Benjamini and Hochberg’s false discovery rate (BH FDR) procedure | | | | | | | | | |

**Table S9 – Post hoc test results for the averaged “r” values of all 8 Corticosterone-Fos correlations using a Welch 1-way ANOVA**

| Games-Howell Post-Hoc Test – Corticosterone and Fos average correlations | | | | | | | | | | | | |
| --- | --- | --- | --- | --- | --- | --- | --- | --- | --- | --- | --- | --- |
|  |  | **0.3REC** | | **0.3REM** | | **1.0REC** | | **1.0REM** | | **COREC** | | **COREM** |
| 0.3REC | Mean difference | — |  | 0.610 | *** | -0.583 | ** | 0.0157 |  | -0.00560 |  | -0.223 |
|  | t-value | — |  | 6.16 |  | -5.44 |  | 0.0917 |  | -0.0347 |  | -0.915 |
|  | df | — |  | 11.0 |  | 12.8 |  | 11.32 |  | 11.85 |  | 8.97 |
|  | p-value | — |  | < .001 |  | 0.001 |  | 1.000 |  | 1.000 |  | 0.933 |
| 0.3REM | Mean difference |  |  | — |  | -1.193 | *** | -0.5942 | * | -0.61542 | * | -0.833 |
|  | t-value |  |  | — |  | -14.98 |  | -38.368 |  | -42.562 |  | -3.572 |
|  | df |  |  | — |  | 13.1 |  | 8.48 |  | 8.72 |  | 7.62 |
|  | p-value |  |  | — |  | < .001 |  | 0.036 |  | 0.020 |  | 0.059 |
| 1.0REC | Mean difference |  |  |  |  | — |  | 0.5988 | * | 0.57752 | * | 0.360 |
|  | t-value |  |  |  |  | — |  | 37.365 |  | 38.411 |  | 1.520 |
|  | df |  |  |  |  | — |  | 9.51 |  | 9.89 |  | 8.07 |
|  | p-value |  |  |  |  | — |  | 0.036 |  | 0.029 |  | 0.663 |
| 1.0REM | Mean difference |  |  |  |  |  |  | — |  | -0.02126 |  | -0.239 |
|  | t-value |  |  |  |  |  |  | — |  | -0.1060 |  | -0.880 |
|  | df |  |  |  |  |  |  | — |  | 13.92 |  | 11.97 |
|  | p-value |  |  |  |  |  |  | — |  | 1.000 |  | 0.944 |
| COREC | Mean difference |  |  |  |  |  |  |  |  | — |  | -0.218 |
|  | t-value |  |  |  |  |  |  |  |  | — |  | -0.819 |
|  | df |  |  |  |  |  |  |  |  | — |  | 11.43 |
|  | p-value |  |  |  |  |  |  |  |  | — |  | 0.958 |
| COREM | Mean difference |  |  |  |  |  |  |  |  |  |  | — |
|  | t-value |  |  |  |  |  |  |  |  |  |  | — |
|  | df |  |  |  |  |  |  |  |  |  |  | — |
|  | p-value |  |  |  |  |  |  |  |  |  |  | — |
| *Note.* * p < .05, ** p < .01, *** p < .001 | | | | | | | | | | | | |
